# Two structural switches in HIV-1 capsid regulate capsid curvature and host factor binding

**DOI:** 10.1101/2022.12.02.518879

**Authors:** James C.V. Stacey, Aaron Tan, John M. Lu, Leo C. James, Robert A. Dick, John A.G. Briggs

## Abstract

The mature HIV-1 capsid protects the viral genome and interacts with host proteins to travel from the cell periphery into the nucleus. To achieve this, the capsid protein, CA, constructs conical capsids from a lattice of hexamers and pentamers, and engages in and then relinquishes multiple interactions with cellular proteins in an orchestrated fashion. Cellular host factors including Nup153, CPSF6 and Sec24C engage the same pocket within CA hexamers. How CA assembles pentamers and hexamers of different curvatures, how CA oligomerization states or curvature might modulate host-protein interactions, and how binding of multiple co-factors to a single site is coordinated, all remain to be elucidated. Here, we have resolved the structure of the mature HIV-1 CA pentamer and hexamer from conical CA-IP_6_ polyhedra to high resolution. We have determined structures of hexamers in the context of multiple lattice curvatures and number of pentamer contacts. Comparison of these structures, bound or not to host protein peptides, revealed two structural switches within HIV-1 CA that modulate peptide binding according to CA lattice curvature and whether CA is hexameric or pentameric. These observations suggest that the conical HIV-1 capsid has different host-protein binding properties at different positions on its surface, which may facilitate cell entry and represent an evolutionary advantage of conical morphology.

**Significance statement:** HIV-1 particles contain a characteristic, conical capsid that shields the genome from the cellular immune system and recruits cellular proteins to direct the capsid to the nucleus. The cone forms from hexamers of CA protein, and twelve pentamers that accommodate curvature. We obtained detailed 3D models of pentamers and hexamers at positions on capsid surfaces with different curvatures. We find two places in CA that switch conformation according to the local capsid curvature and whether CA is in a pentamer or hexamer. We also obtained models of CA bound to peptides from cellular proteins. The data show how switches in CA help it form a cone shape, and interact differently with cellular proteins at different positions on the cone surface.

## Introduction

The mature HIV-1 capsid is a conical, fullerene shell that encompasses the viral genome. It is constructed of a lattice of approximately 200 capsid protein (CA) hexamers incorporating CA pentamers at 12 highly curved vertices (1, 2). The capsid serves multiple functions during viral infection. As the vessel in which reverse transcription takes place, it regulates access of cellular dNTPs (3) whilst simultaneously shielding its genetic cargo from detection and degradation by host cellular immunity systems (4-6). The surface of the conical capsid is dense in protein binding sites that mediate interactions with host machinery necessary for transport of the capsid through the cytoplasm into the nucleus (7-9).

CA monomers consist of two alpha-helical domains. The CA N-terminal domain (CA_NTD_), situated on the outer surface of the capsid, stabilises the hexamer via interactions around a six-fold axis. The C-terminal domain (CA_CTD_) engages in the dimeric and trimeric interactions that join hexamers together into a lattice. The small molecule inositol hexakisphosphate (IP_6_) is known to act as an assembly factor for both immature and mature HIV-1 CA lattices (10, 11). In the mature lattice, IP_6_ coordinates R18 in the central CA_NTD_ pore (12), where it greatly increases capsid stability (13). Flexibility in the linker between the two CA domains and in the interactions between domains have been proposed to provide plasticity necessary to allow the hexamer to construct different lattice curvatures at different positions on the conical surface (14, 15) and may also permit formation of both pentamers and hexamers of CA. The precise stability, or instability, of the capsid is critical for correct function. Accordingly, mutations that hyper-stabilise or destabilise the capsid both lead to marked reductions in infectivity (16), implying that the capsid must retain sufficient stability to maintain a protective function, but also uncoat readily enough at the correct moment.

Low-resolution structures of CA hexamers from capsid cores within HIV-1 virions (1) have validated that high-resolution crystal structures of HIV-1 CA hexamers (17) are representative of hexamers in the virion. In contrast, low-resolution structures of the CA pentamer within virions (1) are not consistent with available crystal structures of crosslinked, mutant CA pentamers (18), and the high-resolution structure of the “in virus” pentamer remains unknown.

Two protein binding sites on the surface of the core have been characterised structurally by way of cryo-EM and x-ray crystallography. The CypA binding loop constitutes a flexible span of residues between helices 4 and 5, and binds cyclophillinA (CypA) (19) and nuclear pore component Nup358, which contains a CypA domain (20, 21). A second binding site within the CA_NTD_ specifically binds phenylalanine-glycine (FG) motifs (22-24). FG-repeat-containing host factors are diverse in their cellular distribution and function (25) and include: Nup153, a nuclear pore complex component; Sec24c, a member of the COPII complex implicated in intracellular vesicle trafficking; and the nuclear-localised CPSF6 (23, 24). All three proteins insert an FG motif into a pocket between helices 3 and 4 of the CA_NTD_, while residues proximal to the FG repeat bind to CA residues in a cleft formed between the CA_NTD_ and the CA_CTD_ of the neighbouring CA in the hexamer. Subtle differences in binding affinity between different FG repeats as a result of varying binding conformation may play an important role in transport and nuclear import of the capsid. Accordingly, quantitative fluorescence microscopy of reverse transcription and pre-integration complexes suggest that there is a hand-off from Nup153 at the nuclear pore to CPSF6 within the nucleus (26). Owing to its critical role in orchestrating early phase replication, the FG-binding site has been the target of concerted efforts to identify CA targeting antiretrovirals. Such studies have led to potent drug candidates including PF-3540074 (PF74) (27) and members of the GS-CA family including lenacapavir (28, 29). A difference in the relative orientations of the CA_NTD_ and the CA_CTD_ between hexamers and pentamers in *in situ* structures (1) alters the apparent accessibility of the FG repeat binding site, but in the absence of high-resolution structures of the viral CA pentamer the implications for host-protein binding to the pentamer are not clear. It is also not clear whether changes in CA hexamer curvature can modulate FG-motif binding.

CA can be induced to assemble *in vitro* in buffers containing IP_6_ at physiological salt concentration to form structures containing hexamers and pentamers that closely resemble authentic CA cores (30) (Highland et al, accompanying manuscript). Here, we present high-resolution structures of CA hexamers and pentamers within *in-vitro* assembled conical CA-IP_6_ polyhedra from multiple lattice contexts differing in local curvature and geometry, alone or in the presence of Nup153, CPSF6 or Sec24C-derived peptides. This analysis suggests that the structure and morphology of the core have evolved to preferentially bind different components of the host-cell machinery at different regions of the conical capsid surface. This is achieved using two structural switches within CA that modulate host-factor binding. The first switch alters the FG-repeat binding pocket between the hexamer and pentamer, precluding FG repeat binding to the pentamer completely. The second switch regulates Nup153 affinity to the hexamer in a manner that is sensitive to local capsid curvature.

## Results

### Structures of the mature HIV-1 CA pentamer and hexamer

Purified recombinant CA protein was assembled in the presence of IP_6_ into mature core-like particles (CLPs), as described previously (30) (Highland et al, accompanying manuscript) (**Fig. S1**). The contents of the reaction mixture were used to prepare cryo-EM grids, which were imaged in the electron microscope using standard single particle data collection conditions (methods and **Table S1**). Inspection of micrographs confirmed that the size and morphology of the CLPs were qualitatively similar to capsid morphologies reported in mature viruses by cryo-electron tomography (cryo-ET) (1), and this was confirmed using cryo-ET (Highland et al, accompanying manuscript).

From these data we selected “particles” corresponding to arbitrary regions of the capsid surface and applied single-particle data analysis techniques including 3D-classification (Methods) (**Fig. S1**). We obtained separate reconstructions of the pentamer and of the hexamer to nominal resolutions of 2.9Å and used these to build atomic models (**Fig. 1A, B, Fig. S2**). Both structures match well to those previously determined at low-resolution within virions (**Fig. S3**), and to those independently determined in the accompanying manuscript (Highland et al, accompanying manuscript).

**Figure 1:**
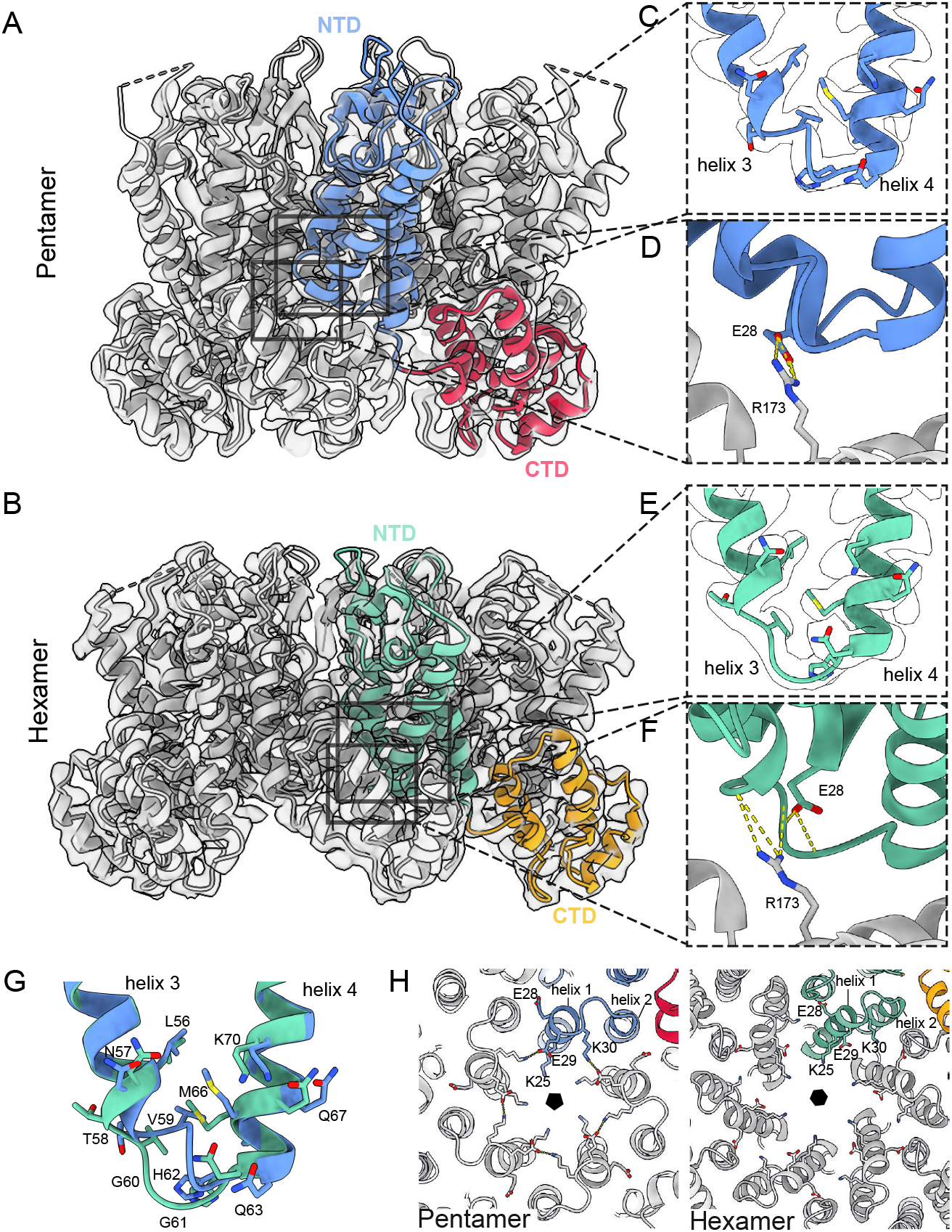
Structure of the HIV-1 CA Hexamer and Pentamer from in-vitro assembled core-like particles. **(A)** Isosurface representation of a single-particle reconstruction of the CA pentamer, viewed from the side. The corresponding atomic model is shown as ribbons in grey, with one monomer coloured in blue (NTD) and red (CTD). **(B)** As in (A), for the hexamer reconstruction from the same data. A single monomer is coloured in green (NTD) and orange (CTD). **(C)** Zoomed in view of the base of helix 3/4 and the intervening loop (FG-repeat binding site) from the pentamer reconstruction and model. **(D)** Zoomed in view of R173, which reaches past the helix 3/4 loop to interact with E28. **(E)** Zoomed in view of the equivalent region of (C), for the hexamer reconstruction and model. **(F)** Zoomed in view of the equivalent region of (D), for the hexamer reconstruction and model. R173 and E28 are separated by the helix 3/4 loop, with which they interact. **(G)** Superposition of models from (C) and (E) reveals structural differences between the pentamer and hexamer at the FG-binding site. As compared to the hexamer (green), the pentamer (blue) has a 3w helical turn at the base of helix 3, V59 is located further towards the centre of the binding pocket and M66 adopts a different conformation. We refer to this region as the hexamer-pentamer switch’. **(H)** View of the central CA_NTD_ pore, from inside the CLP, showing an exchange of binding partners between hexamer and pentamer for charged residues.

The relative orientation of the CA_NTD_ and CA_CTD_ within individual CA monomers is very similar in hexamers and pentamers, with very little change in the position of the inter-domain linker. Comparison of the pentamer and hexamer CA_NTD_ structures revealed structural differences at the base of helix 3 and 4 and the intervening helix 3/4 loop, and includes the formation of an additional 3_10_ helical turn at the base of helix 3 (residues 58-61) in the pentamer (**Fig. 1C-G**). This results in V59 moving towards the core of the CA_NTD_ helical bundle inducing a new conformation of M66. This restructures the pocket which, in the hexamer, is the binding site for FG repeat co-factors Nup153, CPSF6 and Sec24C. Based on comparison with crystal structures of FG repeat peptides bound to CA hexamers, we speculated that in the pentamer this pocket is unable to bind FG motifs. We will refer to this structural rearrangement as the ‘hexamer-pentamer switch’. These observations match and confirm those made in a preprint from the Pornillos lab (Schirra et al 2022, bioRxiv: https://doi.org/10.1101/2022.08.25.505312).

In the hexamer, as in the crystal structure, interactions between neighbouring CA_NTD_s around the symmetry axis are mediated in large part by residues P38 and M39 in helix 2 contacting residues N57 and T58 in helix 3 from the adjacent monomer. In contrast, in the pentamer, helix 2 forms an interface with helix 1 of the adjacent monomer (**Fig. 1H**). Additionally, a new interface forms between helix 1 of adjacent monomers, including a hydrogen bond between T19 and the εN of R18 that may further fix the position of the R18 ring, and a salt bridge between K30 and E29. In the hexamer, K30 is not a pore-facing residue and may interact with E28, while E29 is freely exposed to the centre of the pore (**Fig. 1H**).

Within the central pore of the pentamer we observe two densities, which we interpret as IP_6_ molecules (Highland et al, accompanying manuscript): one IP_6_ sits above, and engages with, the R18 ring, while the other is coordinated by the K25 ring in a fashion that is reminiscent of previously reported hexameric crystal structures (13) (**Fig. S3**). Within the central pore of the hexamer, we also observe two IP_6_ densities co-ordinated by rings of R18 and K25, but density for IP_6_ coordinated by K25 appears to be weaker than that of the pentamer suggesting lower occupancy. The spacing between K25 residues is much larger in the hexamer, and its position suggests that in the absence of IP_6_ it could instead form an intramolecular salt bridge with the exposed E29.

The CA_NTD_–CA_CTD_ interface in the hexamer is essentially identical to that previously described in hexameric CA crystals (PDB: 4XFX; (17)). In the pentamer, a largely hydrophobic interface forms centered around Y169, L211 and M215 in the CA_CTD_, A64, M144, Y145 in the CA_NTD_. R173 in the CA_CTD_, which in the hexamer forms a hydrogen bond with the backbone of N57 and V59 in the adjacent CA_NTD_ (**Fig. 1F**), instead interacts with E28 in the pentamer (**Fig. 1D**).

### Structures of hexamers of varying curvature

To form the surface of the conical HIV-1 capsid, hexamers adapt to different local curvatures at different positions of the core surface, and in certain instances make contact with one or more pentamers. In order to study curvature and contact variation in our sample, and following the approach previously applied to cores within intact HIV-1 virions (1), we first analyzed cryo-ET/subtomogram averaging data of the assembled CLPs (Highland et al, accompanying manuscript), and calculated tilt and twist angles between all pairs of neighbouring hexamers. The distribution of our tilt-twist measurements represents a distribution of curvatures in the CLPs (**Fig. 2**, heatmap) and recapitulates the previous observations made in virions (1). Additionally, through visual inspection of the hexamer-pentamer distributions revealed by cryoET we identified 4 classes of hexamer that make contact with a pentamer: hexamers contacting one (type I), two (2 forms, type II.a and II.b) (**Fig. 2**) and in rare cases three pentamers (type III).

**Figure 2:**
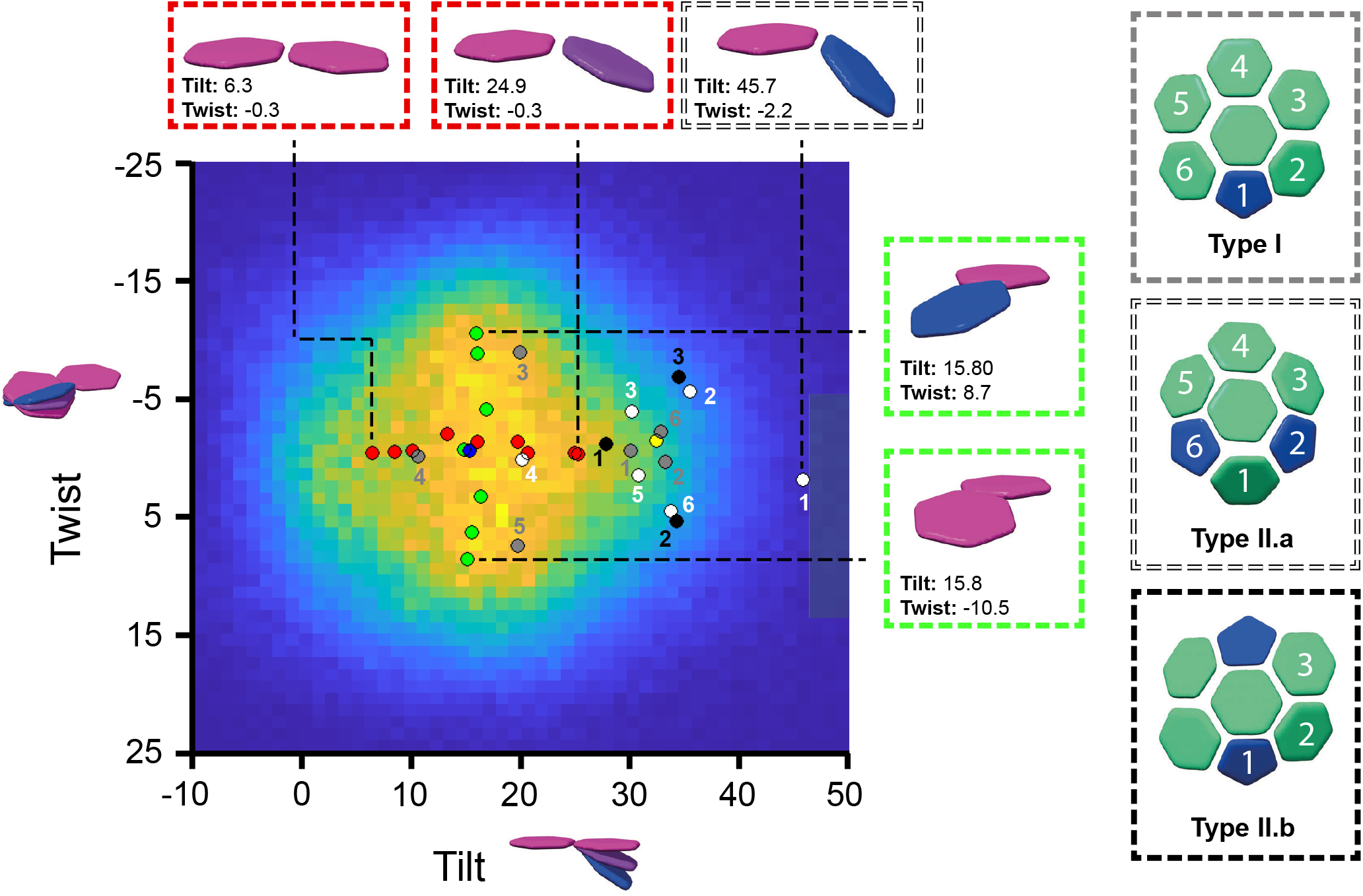
Curvature variation across the mature HIV-1 CA-IP6 cores. Heat map of measured tilt and twist angles for all hexamer-hexamer pairs identified in tomograms. Coloured circles on the heatmap represent the tilt and twist angles of hexamer-hexamer and hexamer-pentamer pairs resolved in single particle reconstructions of individual classes. Blue: the hexamer-hexamer orientation in the average C6 structure from all hexamer data. Yellow: the pentamer-hexamer orientation in the average C5 structure from all pentamer data. Red: class averages from particles grouped according to tilt. Green: class averages from particles grouped according to twist. Numbered circles indicate the orientation from the central hexamer to the adjacent oligomer for hexamers that are immediately adjacent to pentamers, as numbered in the side panels for geometry type I (grey), type ll.a (white), type ll.b (black) − see text for further details.

To study the structures of hexamer curvature variants within our single particle dataset, we applied symmetry expansion to generate a dataset corresponding to all hexamer-hexamer pairs, and then performed 3D variability analysis. From this analysis we derived two primary variability components, which upon inspection resembled hexamer-hexamer tilt in the first class and hexamer-hexamer twist in the second class. We grouped the refined particles along these two variability components to generate nine non-overlapping classes for tilt and seven for twist, each of which was independently refined and reconstructed. Doing so allowed us to generate two series of reconstructions varying by tilt and twist, where all individual reconstructions were at resolutions between 3.1 – 3.4Å. The tilt and twist values for each reconstruction were measured from the symmetry axes of fitted models and mapped onto the distribution determined by cryo-ET (**Fig. 2**, red and green points). Doing so confirmed that the two primary variability components correspond to tilt and twist and that the angular ranges represented in our reconstructions (Tilt: +6.4° - +24.1°, Twist: -10.3° - +9.1°) constitute a significant portion of the true accessible range.

Next, using reference-based classification, we identified classes of hexamer that were adjacent to pentamers within our original hexamer particle set, which we then independently refined. These yielded reconstructions of hexamers contacting a single pentamer, resolved to 3.0Å and two reconstructions of hexamers contacting two pentamers (type II.a and II.b), resolved to 3.6Å and 3.9Å respectively. We were unable to recover a class corresponding to hexamers contacting three pentamers, likely due to its rarity within our sample. To describe the geometry of hexamers adjacent to pentamers we calculated the tilt and twist of all the resolved hexamer-pentamer and hexamer-hexamer pairs. Hexamer-hexamer pairs adjacent to pentamers are highly tilted (**Fig. 2**, black, grey and white points): for example, the two symmetry-related hexamers in the type II.a reconstruction are tilted relative to one another by 45.7°.

### Interpretation of hexamer variant structures and interfaces

The above approach generated 19 reconstructions representing the curvature variability of capsid on the surface of the CLPs. We built models into these reconstructions and compared them with one another. All hexamers, regardless of tilt or twist, share the same structure in the ‘pentamer-hexamer switch’ region (**Fig. 3A**). From this observation we conclude that the structural rearrangement does not represent the most-tilted end of a continuum of increasing lattice curvature, but is a true pentamer-hexamer switch.

**Figure 3:**
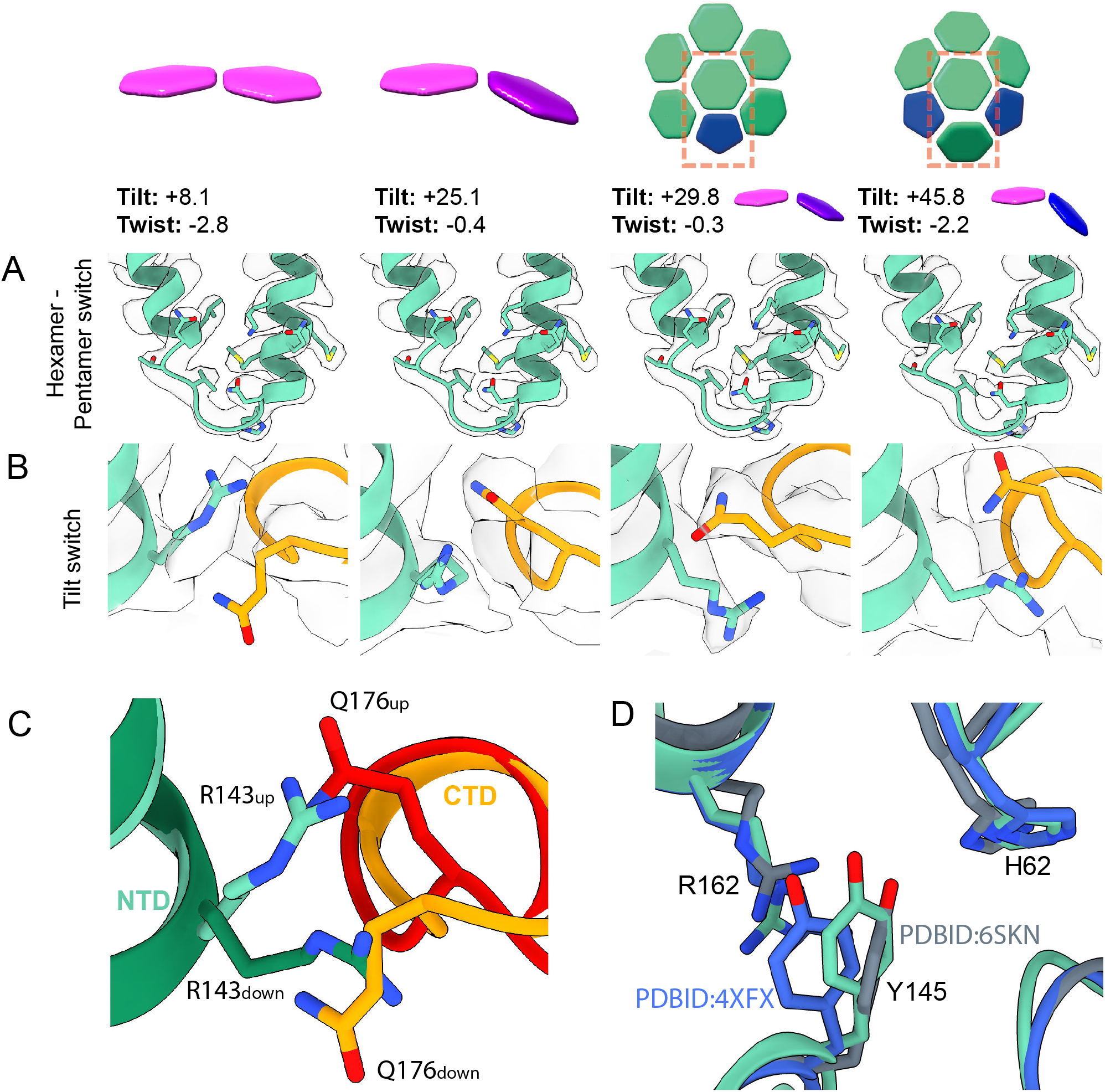
Structures of hexamer tilt variants. **(A)** Isosurface and fit models (green) of the hexamer-pentamer switch region across hexamer tilt variants. The structure of the switch is independent of tilt or twist. **(B)** Isosurface and fit models of the tilt-switch region across hexamer tilt variants. At low tilts, Q176 is below R143, whereas at high tilts Q176 is above R143. At high tilts there is a shift in the backbone in the vicinity of residue Q176, which is not observed in CA monomers forming hexamer-pentamer contacts. **(C)** Superposition of models of the tilt-switch region from CA monomers in low tilt regions (light green (NTD) and light orange (CTD)) and high tilt regions (dark green (NTD) and dark orange (CTD)). **(D)** Superposition of the average hexamer model (green), with PDBID:6SKN (Ni et al., 2020) (grey) and PDBID:4XFX (Gres et al., 2015) (blue), showing the relative configurations of H62 and Y145 as well as R162 of the neighbouring monomer.

We observe curvature-dependent changes at the base of helix 7 in the CA_NTD_ and in the helix 8/9 loop in the CA_CTD_ with which helix 7 forms a small interaction interface (**Fig. 3B,C**). In the average hexamer structure, the R143 sidechain is placed above the helix 8/9 loop, with Q176 pointing downwards. This is the conformation observed in all lower tilt structures (and in the average structure shown in **Fig. 5E**). In contrast, at the most highly tilted hexamer-hexamer contacts, including those in the vicinity of pentamers, R143 is positioned below the helix 8/9 loop and Q176 points upwards. At these positions there is also a shift in the backbone at E175-Q176. At medium tilts we observe a mixture of the two conformations.

R143/Q176 do not appear to reconfigure in response to twist. CA molecules that form hexamer-pentamer contacts form a defined intermediate configuration in which Q176 is above R143, but the backbone at E175-Q176 is in the “low-tilt” conformation (**Fig. 3B**, third column). We will refer to these structural rearrangements as the ‘tilt switch’.

Previous structures of CA hexamers assembled into helical tubes have suggested that Y145 engages in an intramolecular hydrogen bond with H62 in some positions within highly curved hexamers (PDBID: 6SKN), contrasting with the situation in planar hexamer crystals where Y145 forms an intermolecular contact with R162 (PDBID: 4XFX) (14, 17, 31). In contrast, we observe no curvature dependence of interactions involving Y145, H62 and R162 in our curvature variant structures – in all cases, as in the average hexamer structure, the position of Y145 is intermediate between the two previously observed positions (**Fig. 3D**). Indeed, the entire inter-domain hinge, of which Y145 is the N-terminal residue, is very similar in all structures (**Fig. 4A**).

**Figure 4:**
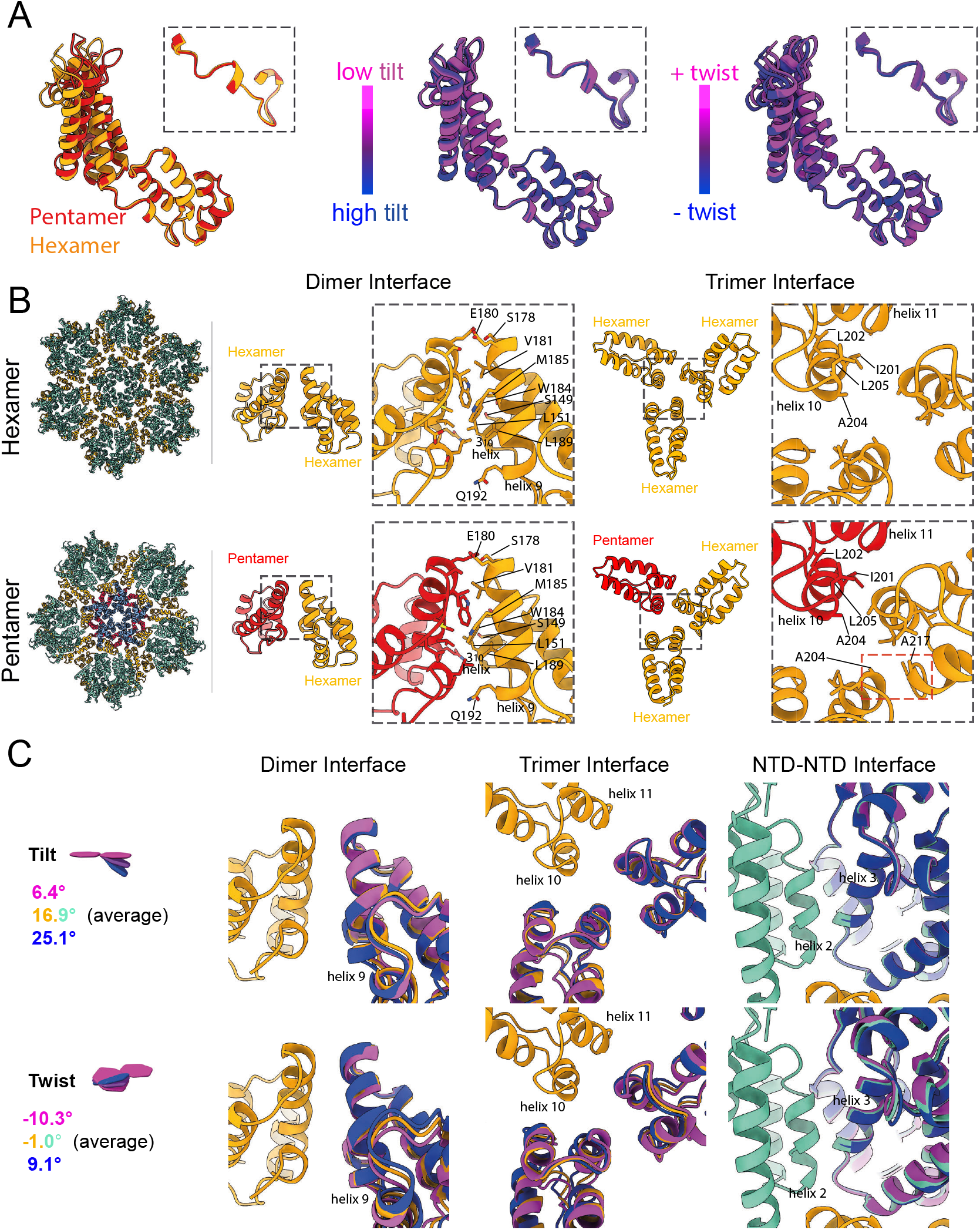
Flexibility of CA and its interactions. **(A)** Superposition of CA structures from the hexamer (orange), and pentamer (red). Superpositions of structures obtained from different tilt/twist classes, coloured blue to magenta. Structures are aligned on the CA_CTD_, insets illustrate that there is minimal motion around the inter-domain hinge. (**B)** The dimeric interface between CA molecules mediated by helix 9, and the trimeric interface mediated by helix 10, for interfaces involving only hexamers (orange), or also including pentamers (red). The dimeric interface is largely conserved, whereas trimeric contact points including a pentamer form a new interaction between helix 10 and 11. **(C)** Superposition of structures obtained from different tilt/twist classes at the dimeric and trimeric interfaces, as well as in the region of helices 2 and 3 in the NTD. Minimal structural changes are observed at the dimeric interface, whereas the trimeric interface and interface between NTDs show rotations at hydrophobic interfaces.

We next analysed the protein-protein interfaces that mediate hexamer-hexamer interactions at the dimeric and trimeric interfaces in the lattice, as well as at the quasi-equivalent interfaces involving the pentamer. The dimeric interface formed by helix 9 is remarkably invariant across the core surface, and matches the structure and minimal plasticity observed in previous studies (14, 17) (**Fig. 4B**). This is in contrast to earlier suggestions that variable curvature is accommodated based on plasticity of the dimerization interface (15). The three-fold lattice interface formed by helix 10, however, shows variation. At interfaces between three hexamers (independent of tilt and twist angle), helix 10 engages in a symmetrical three-helix bundle, held in place by hydrophobic patch which includes residues I201, L202, A204 and L205, as previously observed in helical arrays (15). At interfaces between a pentamer and two hexamers, one hexamer helix is removed from the bundle and instead forms a new contact with helix 11 of the other hexamer, involving residues A204 and A217 (**Fig. 4B**). There is a small amount of flexibility in the precise structures of hexamer-hexamer three-fold lattice interfaces as a result of tilt and twist variation (**Fig. 4C**), likely facilitated by the non-specific nature of the hydrophobic interactions at this site, however none of these structures resemble the structure at the hexamer-pentamer interface. The hydrophobic helix 2-3 interface around the central pore also provides a small degree of flexibility, allowing relative motion of neighbouring CA_NTD_ domains in response to lattice bending and twisting (**Fig. 4C**).

### Structures from cores bound to FG-repeat peptides

To investigate what effect the hexamer-pentamer switch and the tilt switch have on host protein binding we next repeated our structural analysis on cores incubated with FG repeat containing peptides; Nup153_(1407-1429)_, CPSF6_(276-290)_ and Sec24C_(228-242)_. All three of these peptides have been described to bind to mature CA hexamers via an FG motif (23, 24). We determined structures of hexamers from peptide-bound cores to nominal resolutions of 2.6Å, 3.1Å and 3.1Å and associated pentamers to 3.1Å, 3.5Å and 3.2Å respectively (**Fig. S2**). Within all three hexamer reconstructions we observe unambiguous density for the corresponding peptides (**Fig. 5A**) which adopt structures essentially identical to previously reported crystal structures (22-24). In contrast, we do not detect any density corresponding to peptides bound to pentamers (**Fig. 5B**), though in all three cases peptide density is visible in the binding pocket of the hexamers immediately adjacent to the pentamers (**Fig. S4**).

**Figure 5:**
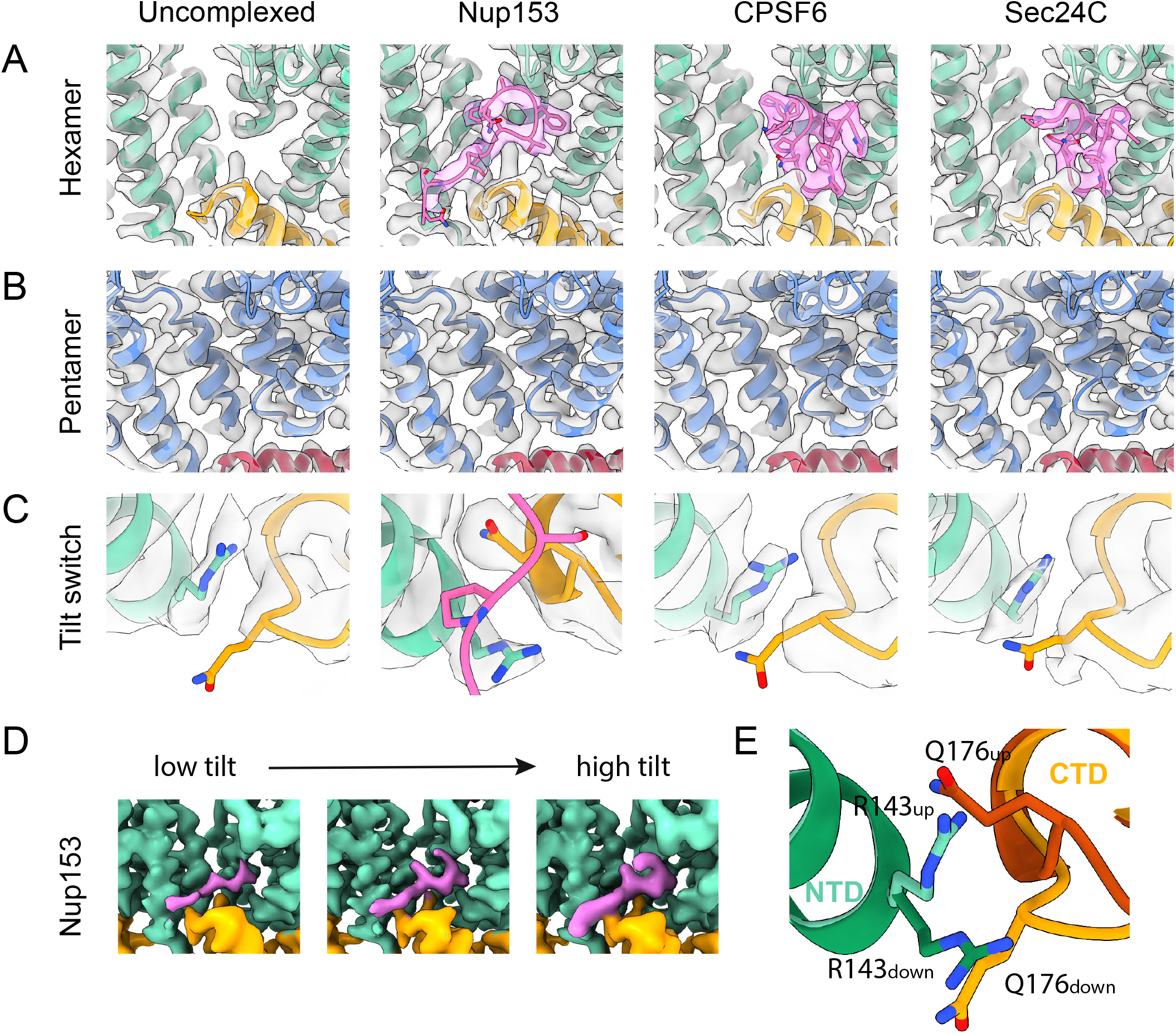
The hexamer-pentamer switch and the tilt switch regulate cofactor binding. **(A)** Isosurface representation and ribbon models of the FG-binding pocket of the CA hexamer reconstructed from cores incubated without peptide and with peptides from Nup153_(1407-1423)_, CPSF6_(313-327)_ and Sec24C_(228-242)_. The NTD is coloured green, the CTD is orange and peptide is pink. **(B)** The equivalent pocket in the pentamer is empty in all four cases. **(C)** Zoomed in view of the tilt-switch region of the hexamer reconstructions for each of the four samples, coloured as in (A). The Nup 153_(228-242)_ bound reconstruction has a tilt-switch conformation the same as observed in highly-tilted uncomplexed samples. **(D)** Isosurface volumes of Nup153_(1407-1423)_ bound CA monomers engaging in low (+6.3°), medium (+16.0°) and high-tilt (+24.9°) lattice interactions, coloured as in (A). Nup153_(1407-1423)_ occupancy increases at higher tilt angles. **(E)** Superposition of the tilt-switch region from the uncomplexed hexamer (light colors) and the Nup153_(1407-1423)_ bound hexamer (dark colors).

From these observations we conclude that the hexamer-pentamer switch described above prevents binding of the FG motif in these peptides to the pentamer. Superposition of our peptide-bound hexamer CA_NTD_ structures and our pentamer CA_NTD_ structure confirms that residue M66 in the pentamer would clash with the binding position of phenylalanine within the pocket (**Fig. S5**).

Further comparison of our peptide bound and unbound hexamer structures revealed that the tilt switch adopts the bulk, lower-tilt conformation in the average hexamer in the CPSF6_(276-290)_ and Sec24C_(228-242)_ bound capsids, as seen in the unbound apo hexamer. In contrast, in the Nup153_(1407-1429)_ bound capsid, the switch is in the high-tilt conformation (**Fig. 5E**) with R143 positioned below the helix 8/9 loop, Q176 pointing upwards and the shift in the backbone position at E175-Q176. The peptide-bound structures are, in all cases, consistent with the available crystal structures, suggesting that the interaction of P1411 and S1412 in Nup153 with Q176, A177 and R143 in CA stabilizes the high-tilt conformation of the tilt-switch (**Fig. 5C**). The FG motifs in Sec24c and CPSF6 adopt a more compact structure and do not interact with these residues of CA. We proceeded to classify the Nup153_(1407-1429)_ in bound hexamers according to tilt/twist and position relative to pentamers to generate context-specific structures exactly as described above for the apo capsid. This analysis yielded structures at resolutions from 3.1 – 3.7 Å. Inspection of these resulting densities revealed no detectable changes in the tilt-switch or in the Nup153 binding pocket, suggesting that Nup153 binding stabilizes the high-tilt conformation of the switch at all tilt angles. We did however, detect an increase in relative Nup153 peptide density at higher tilt angles, suggesting increased peptide occupancy (**Fig. 5D**). We posit that the increase in observed Nup153_(1407-1429)_ occupancy at higher tilts correlates with the arrangement of the R143/Q176 tilt switch in the apo form, because the high-tilt conformation is favourable for Nup153_(1407-1429)_ binding.

## Discussion

CA must be flexible enough to accommodate curvature differences across the surface of the conical core, where angles between neighboring hexamers can vary by 30°. It must also be flexible enough to fill both pentameric and hexameric positions in the core surface. The scenario once considered most likely was that flexibility was accommodated by the inter-domain hinge and the dimeric interface involving helix 9, both of which vary in solution and in helical arrays of CA (14, 15). Our previous low-resolution analysis of the CA flexibility within intact virions suggested that, instead, flexibility is accommodated by small structural changes distributed throughout CA (1), and this is largely consistent with recent analysis of helical CA arrays (14). The higher-resolution data presented here allow structural changes to be analyzed at the amino-acid level. Both the inter-domain hinge and the dimeric interface are homogeneous across the full range of hexamer-hexamer curvatures, and at interfaces between hexamers and pentamers. As suggested by the low-resolution analysis, variable curvature is indeed accommodated by small, distributed changes that do not impact the local bonding interactions between amino acids, but that can combine over longer distances to change curvature. Hydrophobic interfaces at the three-fold axis and in the helix 2-3 interface around the central hexamer pore provide slippery surfaces that can help accommodate flexibility of the hexamer. Within the hexameric lattice the only clear exception to this general observation is the tilt switch.

How does CA adapt its structure at the pentameric positions in the lattice? The 2-fold interface at helix 9 formed by the pentamer-forming CA_CTD_ with the neighboring hexamer is essentially the same as the interface between two hexamers, while the hydrophobic interfaces at helix 10/11 adapt to the change in geometry at the pseudo-three-fold axis between one pentamer and two hexamers. The relative positioning of the CA_NTD_ and CA_CTD_ within the monomer is also very similar between pentamers and hexamers. Indeed, superimposing the CA_CTD_ of a monomer from the hexamer, with that of a CA monomer from the pentamer, results in a closely overlapping CA_NTD_ with minimal difference in the hinge orientation (**Fig. S6**). Overall, the impression is that the CA lattice around the five-fold position continues to grow via a conserved two-fold interface into the five-fold position. This growth positions the CA_NTD_ relative to the neighboring CA_CTD_ such that it would lead to a clash of the CA_NTD_ loop between helices 3 and 4, and R173 in the neighboring CA_CTD_ (**Fig. S6**). This clash is resolved by the ‘hexamer-pentamer switch’ which alters the position of this loop, allowing interaction between E28 and R173 to stabilize the pentamer-specific packing of CA_NTD_ and neighboring CA_CTD_ (**Fig. 1D**). This may lead to an exchange of interactions: in the pentamer E29 interacts with K30, while in the hexamer E28 may interact with K30, leaving E29 exposed in the pore to possibly interact with K25 in the absence of IP6 (**Fig. 1G**). The E28A/E29A double mutant, as well as the R173K mutant, are able to assemble and release immature-like particles but are non-infectious (32, 33). The pentameric packing results in a much more compact arrangement of helix 1 around the pore and a much greater charge density where the K25 residues from five CA molecules are close together. This provides a possible explanation for why pentamer formation, in contrast to hexamer formation, is dependent on K25 and IP6 or similar charged molecules for assembly (Highland et al, accompanying manuscript).

The hexamer-pentamer switch also alters the conformation of the FG repeat binding pocket. Our structures in the presence of host peptides show that this change prevents binding of FG repeat peptides from Nup153, CPSF6 and Sec24c to this pocket in the pentamer. What are the possible implications of this for core function? On one hand it lowers the density of FG repeat binding sites at the tips of the core, in particular at the highly-curved tip of the narrow end of the core where pentamers are enriched. This may facilitate access for proteins that bind elsewhere on CA, for example the cytoplasmic Nup358 may more easily interact with the CypA loop at the tips of the cores if the occupancy of other proteins to the FG repeat binding site is lower. On the other hand, it provides a pentamer-specific structure, that may provide a binding site for as-yet-unknown host proteins that are required for transport to or into the nucleus.

The tilt-switch provides a second mechanism by which host proteins might preferentially bind particular regions of the core surface. It would allow Nup153 to preferentially bind to the more curved regions of the core which are localized towards the ends of the cone. Where different cellular proteins are competing for the same pocket in capsid, this might in turn lead to other proteins binding to flatter parts of the capsid surface What are the possible implications of this for core function? On one hand, it may allow different host proteins to cluster on different regions of the core surface, creating local functional surfaces with high allostery. On the other hand, it may orient the core in defined ways, for example positioning regions of higher curvature towards regions of high Nup153 density during nuclear entry (34).

We speculate that maintaining local regions of core surface that specialize in interaction with different host proteins is one of the evolutionary advantages provided by the unusual conical shape of the HIV-1 core.

## Acknowledgements

This work was funded by: National Institute of Allergy and Infectious Diseases under awards R01AI147890, HIVE-2 Collaborative Development Program 5U54AI150472-09, and U54 AI170855-01 to R.A.D, the UK Medical Research Council MC_UP_1201/16 to J.A.G.B. and the Max Planck Society to J.A.G.B.. Single particle data was collected at MPI Biochemistry. We thank Dustin Morado and Zunlong Ke for assistance with data collection; Dominik Hrebik and Hui Guo for advice on data analysis; Florian Beck for assistance with computing infrastructure; Carolyn Highland for comments on the manuscript.

## Author contributions

J.C.V.S., R.A.D. and J.A.G.B. designed research; J.C.V.S., A.T. and R.A.D. prepared samples; A.T. and J.L. performed preliminary experiments. L.C.J. provided reagents and guidance. J.C.V.S. carried out SPA cryo-EM and related data processing; J.C.V.S. and J.A.G.B. analyzed and interpreted structural data; J.C.V.S. prepared the figures; J.C.V.S. and J.A.G.B. wrote the manuscript with input from all authors; R.A.D. and J.A.G.B obtained funding and managed the project.

## Data Sharing

Structures determined by electron microscopy are deposited in the Electron Microscopy Data Bank under accession codes EMD-XXXXX – EMD-XXXXX. Corresponding molecular models are deposited in the Protein Data Bank under accession codes XXXX – XXXX. Any additional information required to evaluate the conclusions of the paper is included in the paper or available from the lead author on request.

## Materials and methods

### Sample preparation

Protein expression and purification, and in vitro assembly of CLPs were performed exactly as described in the accompanying manuscript (Highland et al, accompanying manuscript).

### Cryo-EM grid preparation

C-Flat CF-2/2-3Cu-50 grids were glow-discharged for 45 seconds with a current of 25 mA in a PELCO easiGlow glow discharger immediately before use. All samples were vitrified using a Thermo Fisher Scientific Vitrobot Mark IV, operated at 100% humidity and 18°C.

In the case where no peptide was added to the conical CA-IP_6_ cores, the sample mixture was made by mixing CLPs in assembly buffer with BSA-conjugated 10 nm gold fiducials in 1× PBS at a ratio of 8:1. For peptide binding experiments, an additional peptide-binding step was carried out in order to prepare the sample for plunge-freezing. Conical CA-IP_6_ cores were mixed with a volume of BSA-conjugated 10 nm gold fiducials in 1× PBS, calculated to give a final core:gold ratio of 8:1 after peptide addition. The peptides used were Sec24C residues 228-242, CPSF6 residues 313-327 and Nup153 residues 1407-1423 and were obtained either from Donna Mallery (MRC Laboratory of Molecular Biology, Cambridge, United Kingdom) (synthesised by Designer Biosciences) (Nup153 and CPSF6) or the Max-Planck Institute for Biochemistry Peptide Service (Sec24C) as a lyophilized powder. A solution of the respective peptide at 10 times the required concentration, in core assembly buffer containing 1 % DMSO, was then diluted 1:10 in this mixture of cores and gold fiducials, and incubated on ice for 15 minutes prior to use.

4μl of the sample mixture for plunging was applied to the carbon side of grids within the humidity chamber of the Vitrobot. The sample was then manually blotted from the opposite side of the grid for 3 seconds using Whatman No. 1 filter paper, and then plunge-frozen in liquid ethane. Grids generated in such a way were compatible with both tomography and single particle data acquisitions.

### Single particle data collection

All data for single particle analysis were collected on a Titan Krios G3i cryo-Electron TEM (TFS) operated at 300 keV equipped with a Falcon 4 direct electron detector. Movies were collected at a nominal magnification of x130,000 with a resulting pixel size of 0.93 Å and a total accumulated dose of ∼40 e^-^/ Å at under focus values ranging from -0.6 to -3.0 μm in steps of 0.2 μm. Data were collected as movies automatically with EPU acquisition software (TFS). Data collection parameters are summarised in Extended Data Table 1.

### Image processing

Dose-fractionated movies were aligned, dose-weighted and averaged with MotionCor2 (35) in RELION-4.0 (36). Automated particle picking was performed using crYOLO (37) motion corrected micrographs. Initial pick annotations and model training was performed independently for each of the four single-particle datasets. In order to generate an initial annotated training set, a subset of 100 micrographs of each dataset were manually picked in the crYOLO boxmamanger GUI; overlapping picks were placed across the entire visible surfaces of HIV-IP_6_ cores without attempting to define individual hexamers or pentamers, whether such features were visible or not.

Motion corrected micrographs and picked positions were imported into cryoSPARC (38, 39) where initial defocus estimates were calculated using Patch CTF estimation. Using the same initial set of picked particles, either refined hexamer or refined pentamer positions were derived with parallel rounds of heterogeneous refinement using either hexamer or pentamer reconstructions as initial references, both of which were obtained previously from subtomogram averaging (Highland et al, accompanying manuscript). Resulting classes from these refinements that did not resemble the targeted structure, or were of visibility low quality, were discarded and particles from selected classes were pooled and used for further refinement.

For structures of hexamers adjacent to pentamers, particle positions were derived in one of two ways; either through 3D-classification of previously derived hexamer positions, as was the case for the uncomplexed and Nup153_(1407-1423)_ datasets, or through symmetry expanding pentamer positions and shifting of the particle box onto the neighbouring hexamer positions, as was the case for CPSF6_(313-327)_ and Sec24C_(228-242)_ bound datasets. Further 3D-classification of these particles was employed to detect further type II.a and II.b arrangements.

In all cases, once initial particle positions were identified, non-uniform refinement was performed, followed by local ctf-refinement and then local masked-refinement with updated CTF values. Where an increase in resolution could be gained, additional rounds of heterogeneous refinement followed by local refinement were performed. For the uncomplexed and Nup153_(1407-1423_)-bound datasets, particle positions were imported back into RELION-4.0, using pyem, where particle motions were corrected using Bayesian polishing. Polished particles were then imported back into cryoSPARC where a final round of local refinement was performed.

#### Building and refinement of atomic models

All CA models were derived principally from a crystal structure of full-length HIV-1 CA, PDB: 4XFX (17). Initial coordinates of peptide ligands were sourced from PDB:6PU1 (Sec24C), PDB:5STX (Nup153) and PDB:4U0A (CPSF6). The CA_NTD_(1-147), CA_NTD_(148-230) and peptides were independently docked into their respective density as rigid bodies within UCSF chimera (40). Atomic positions and geometry were refined using ISOLDE as a plugin within UCSF ChimeraX (41, 42). Manual adjustment of side chain rotamers was performed in COOT. For non-symmetrical hexamers adjacent to pentamers this was performed for all six CA chains independently. Models were finally refined as complete hexameric or pentameric assemblies using PHENIX.real_space_refinement (43) with non-crystallographic symmetry enforced where appropriate. Validation statistics were calculated using MolProbity (44). A summary of model validity can be found in Extended Data Table 2.

The above approach was used to generate models for Apo and peptide-bound average pentamers, average hexamers and Type 1 hexamers next to pentamers. For Type 2 hexamers next to pentamers and for tilt and twist classes, domains from monomers from the average structures were fitted as rigid bodies and local regions relevant for interpretation were manually adjusted.

### Tilt/twist analysis of CLPs

Cryo-ET data, as well as the methods for data collection and processing, is described in the accompanying manuscript. The tilt and twist distribution (heatmap in **Fig. 2**) was calculated from subtomogram positions and orientations exactly as described in Mattei et al., 2016.

### Tilt/Twist classification of CA-IP_6_ hexamer-hexamer from single particle reconstructions

In order to derive tilt and twist classes for hexamer-hexamer pairs our final C6 hexamer reconstruction was subjected to symmetry expansion and then 3D variability analysis in cryoSPARC (45). The mask used for 3D variability analysis consisted of the central hexamer and one neighbouring hexamer. 3D variability components corresponding to tilt and twist could be identified by visual inspection, and consistently were found within the top 3 reported components. Particles were grouped along the two variability components, 9 groupings for tilt and 7 for twist. Pooled particles were then subjected to local refinement with no symmetry applied to yield a final reconstruction.

### Measuring tilt/twist of single particle reconstructions

Previously deposited hexameric (5MCX) or pentameric CA structure (5MCY) (1), models were rigidly fit into adjacent densities using UCSF Chimera. Within each of the two fit models, two positions were defined along the symmetry axis using the ‘structure measurements’ feature within UCSF Chimera. From the defined centroid positions, vectors along the symmetry axis of both oligomeric structures were derived. Tilt/twist angles were then calculated in the same way as described previously (1), and were compared to the tilt-twist distribution measured from cryo-ET data. Cryo-ET data is described in the accompanying manuscript (Highland et al, accompanying manuscript), and the tilt-analysis of the data was performed exactly as described in (1).

---

**Figure S1a:**
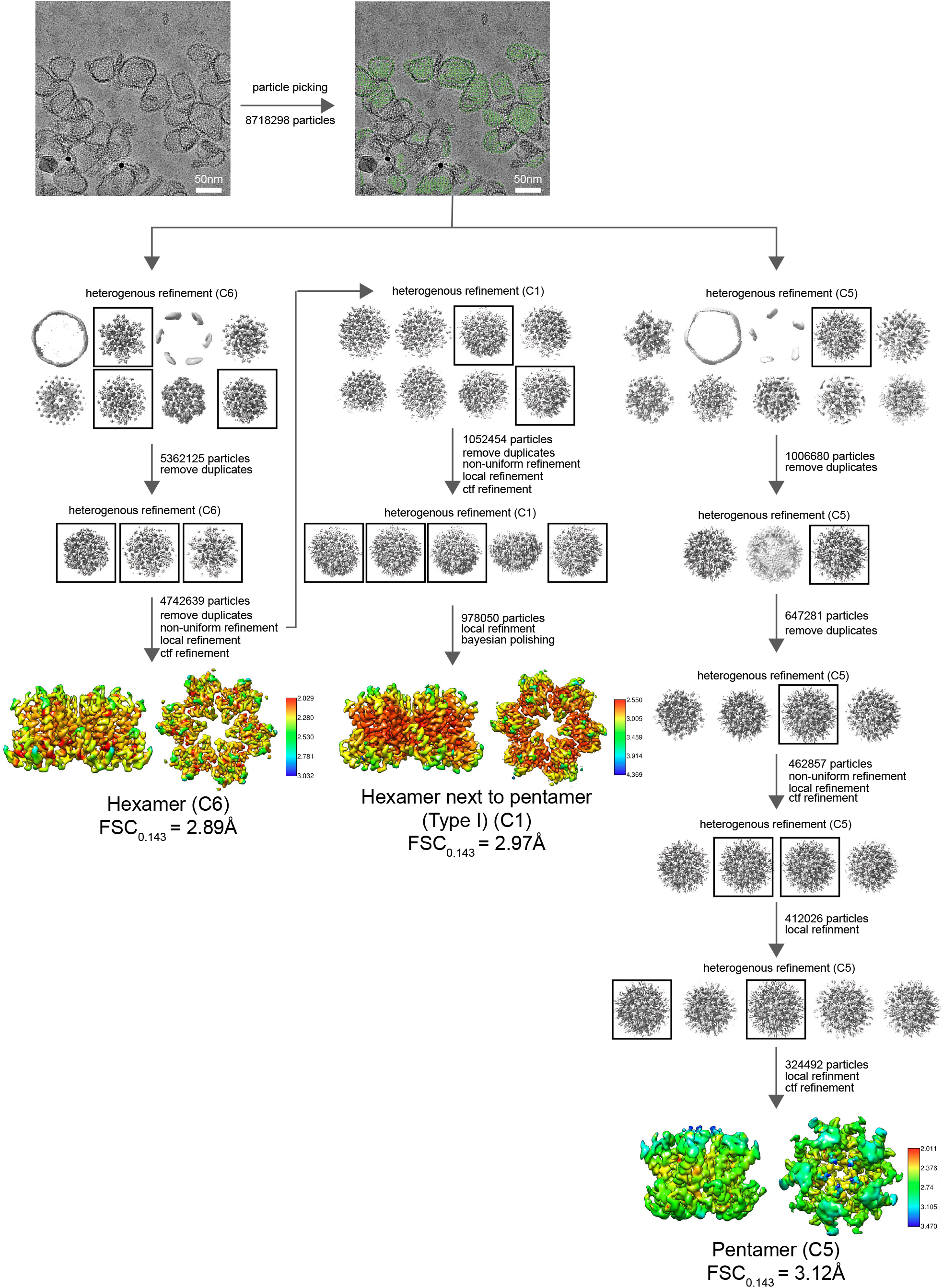
Workflow for picking and processing of cryo-EM data: Apo Structures

**Figure S1b:**
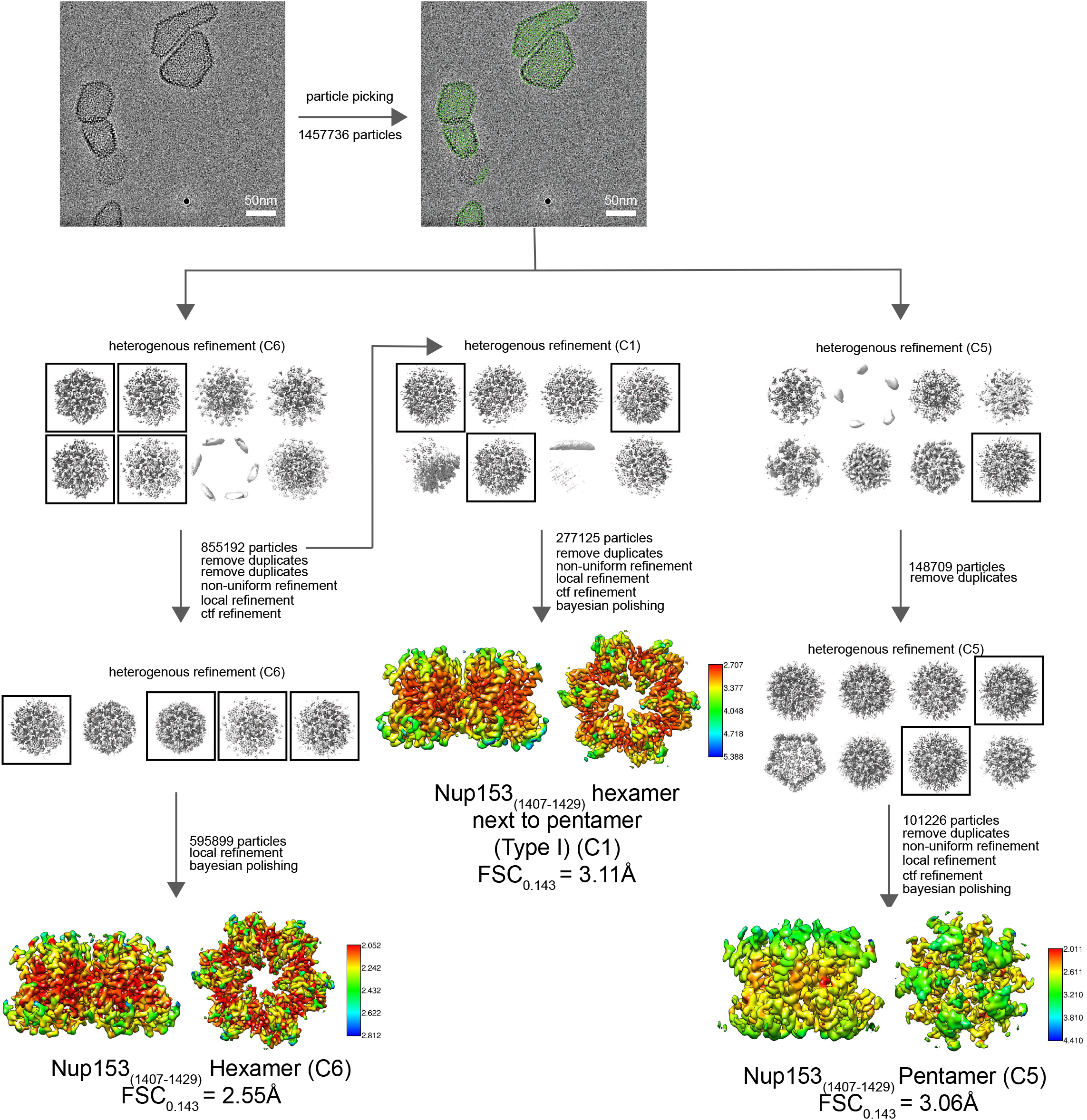
Workflow for picking and processing of cryo-EM data: Nup153 Structures

**Figure S1c:**
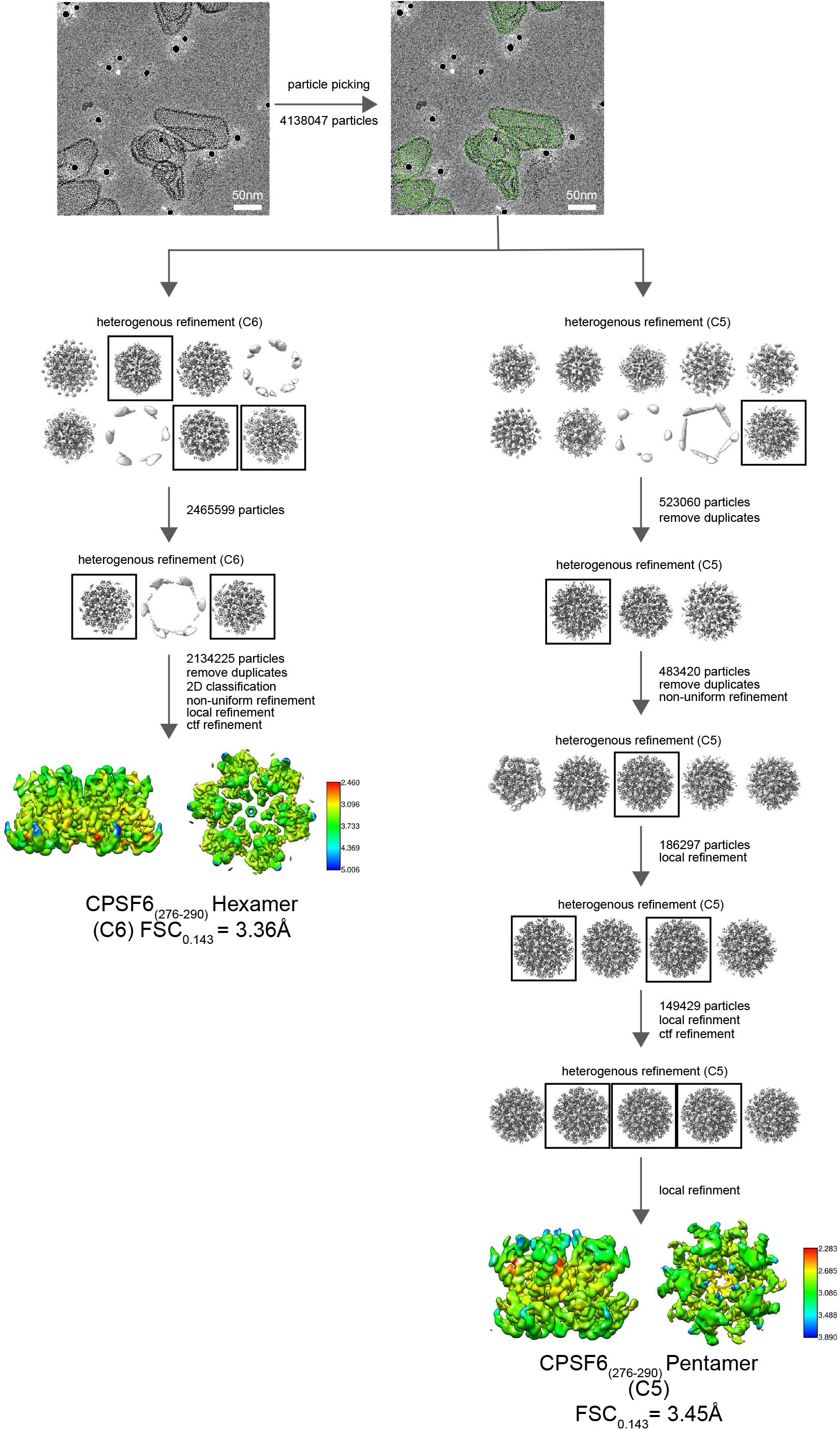
Workflow for picking and processing of cryo-EM data: CPSF6 Structures

**Figure S1d:**
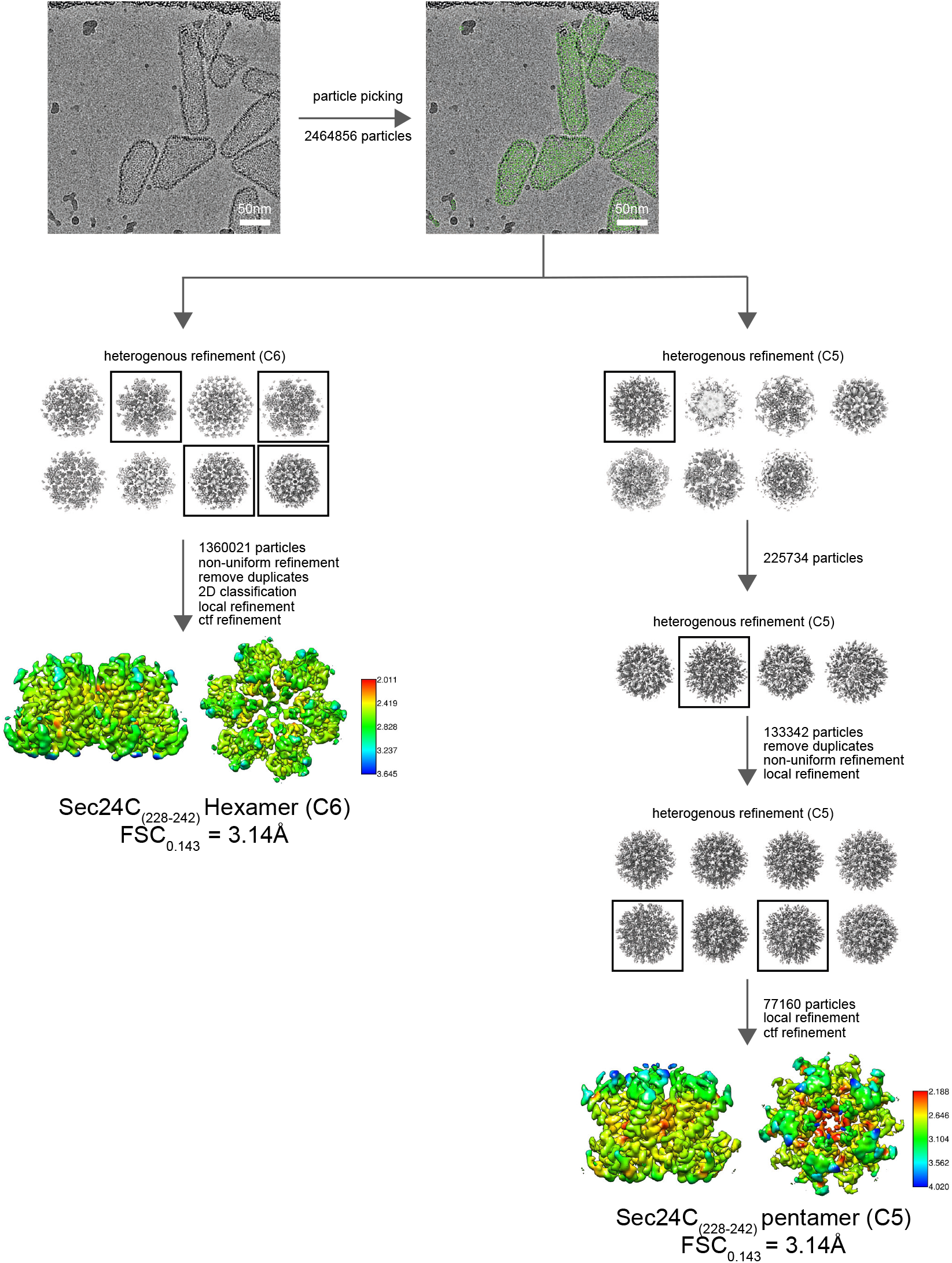
Workflow for picking and processing of cryo-EM data: Sec24C Structures Pipeline for picking and processing of cryo-EM images of CLPs bound to no peptide (Apo), Nup153_(1407-1429)_. CPSF6_(276-290)_ or Sec24C_(228-242)_. After automated picking of CLP surfaces in crYOLO, heterogenous refinement was used to sort hexamer and pentamer position as well to identify poor quality particles, which were discarded. In the case of the Apo and Nup153_(1407-1429)_ bound datasets, further classification was performed on the hexamer positions to identify hexamers directly adjacent to pentamers. Local resolution maps of all the final reconstructions, calculated in cryoSPARC, are shown.

**Figure S2:**
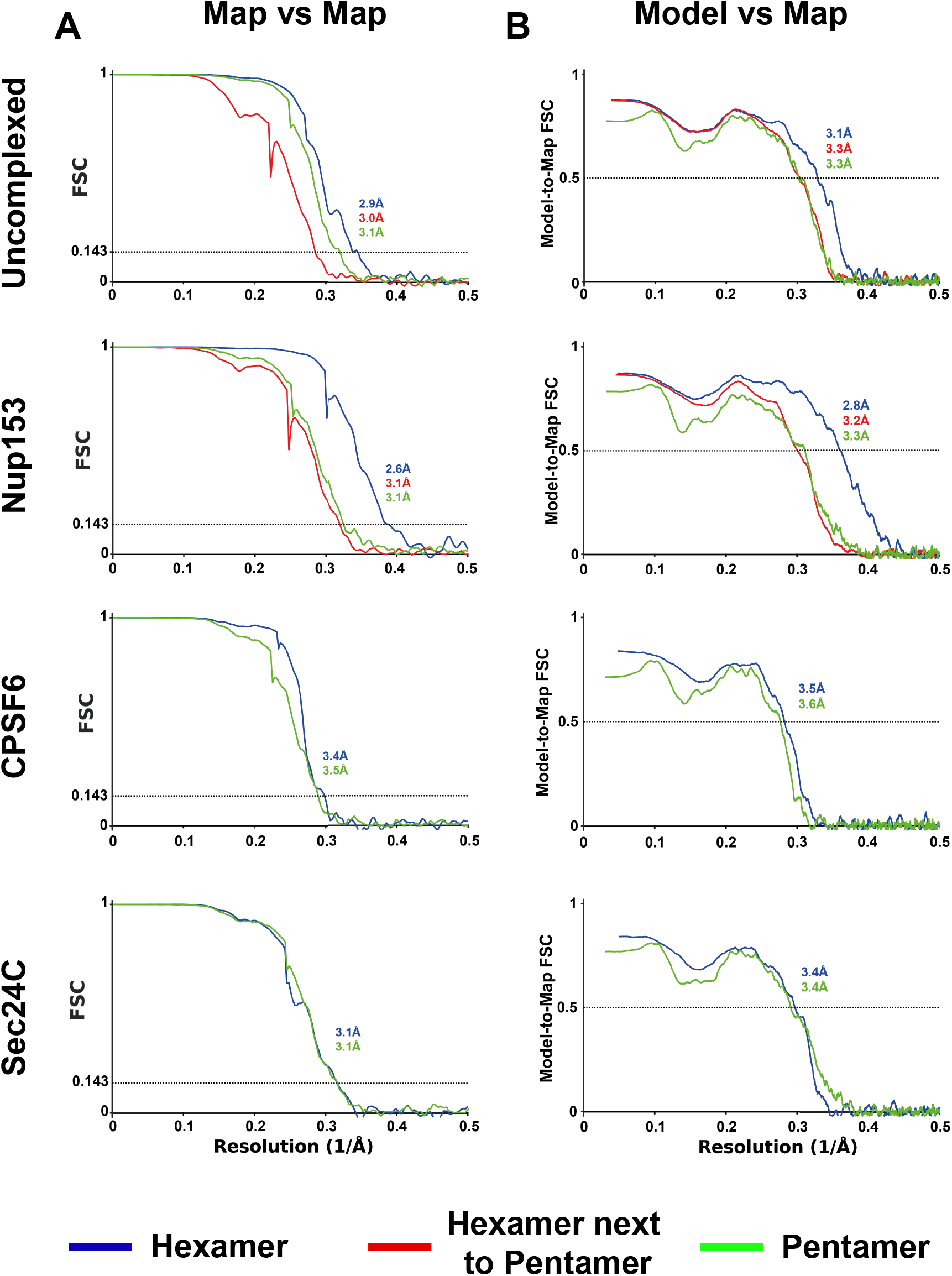
Resolution assessment of cryo-EM structures. **(A)** Global resolution assessment of final reconstructions by Fourier shell correlation for each of the four datasets. **(B)** Fourier shell correlation between respective models and maps.

**Figure S3:**
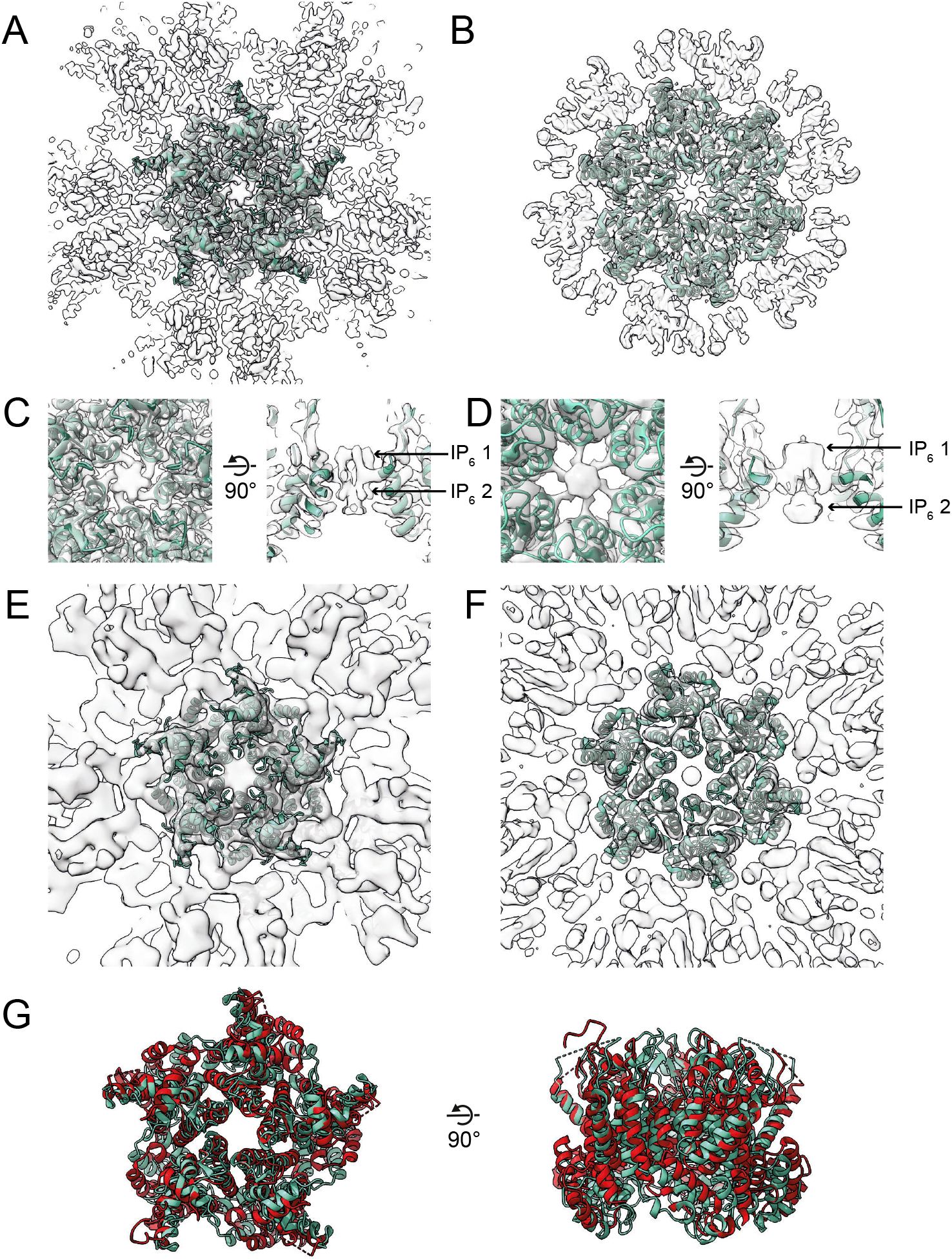
Comparison of hexamer and pentamer structures with in-virus structures. **(A)** Model of the HIV-1 CA pentamer (green) determined from non-peptide bound CLPs is fit into the corresponding density (grey). **(B)** As in (A), for the hexamer. **(C)** Zoomed in view of the central pore region from the top (left) and side in cross-section (right), showing density that corresponds to two IP_6_ molecules. **(D)** As in (C), for the hexamer, also showing two densities that correspond to IP_6_. **(E)** Model from (A and C), fit into a reconstruction of the HIV-1 pentamer determined from intact virus particles by subtomogram averaging (EMD:3466) (Mattei et al., 2016). **(F)** Model from (B and D), fit into a reconstruction of the HIV-1 hexamer determined from intact virus particles by subtomogram averaging (EMD-3465) (Mattei et al., 2016). **(G)** Superimposition of the model from (A and C) with a previous pentamer structure engineered by the addition of disulfides and determined by X-ray crystallography (PDBID:3P05, red) (Pornillos et al., 2011), the structures are distinct.

**Figure S4:**
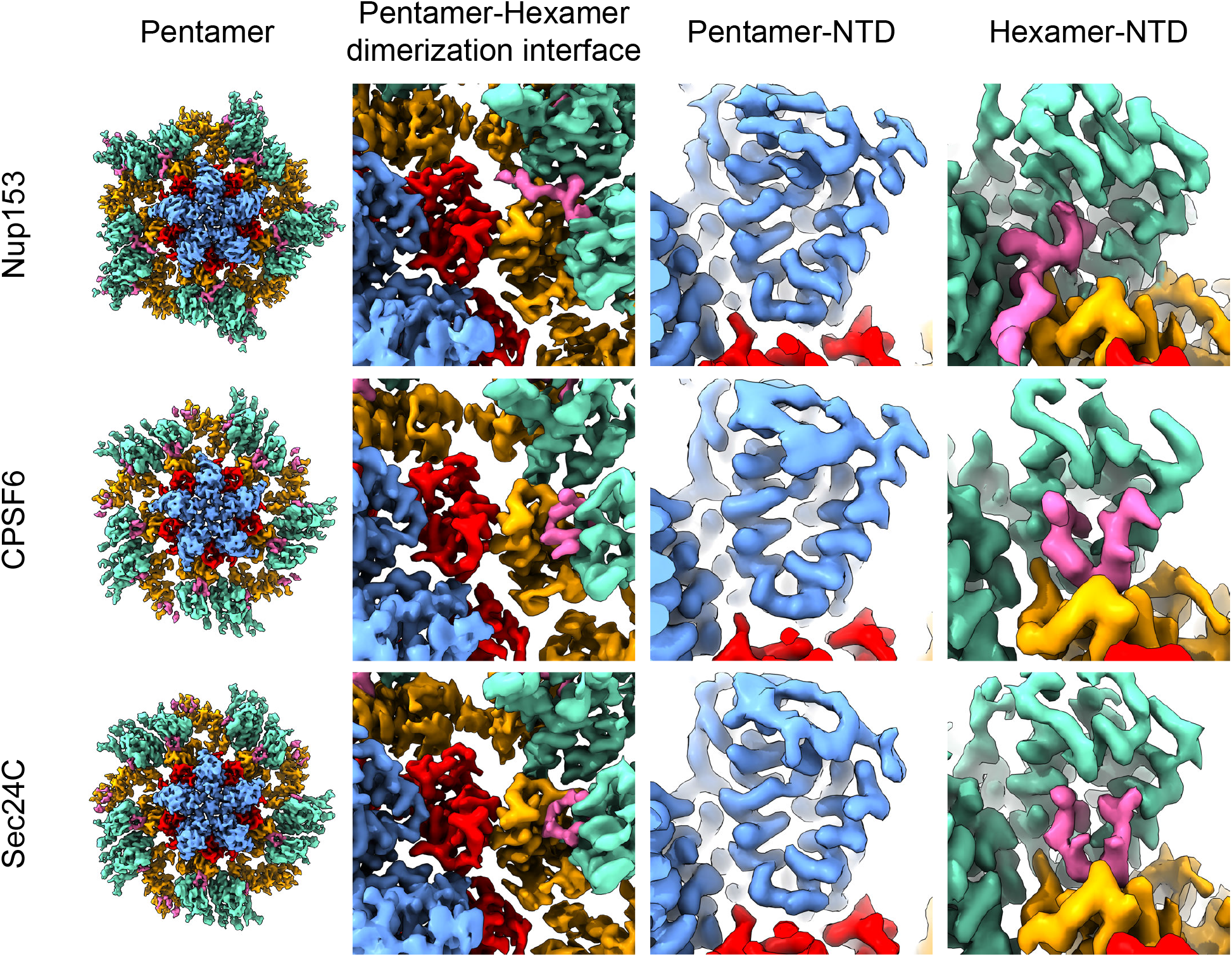
Peptide density observed in hexamers adjacent to pentamers. Reconstructions of the HIV-1 CA pentamer determined from CLPs incubated with Nup153_(1407-1429)_, CPSF6_(276-290)_ or Sec24C_(228-242)_. Density corresponding to the pentamer NTD and CTD are coloured blue and red respectively. Density corresponding to neighbouring hexamer molecule NTDs and CTDs are also resolved and are coloured green and orange respectively. Density corresponding to bound peptide within reconstructions is coloured pink. The second column shows a zoomed in view of the hexamer-pentamer dimerization interface for corresponding reconstructions. The third column shows a zoomed in view of corresponding pentamer NTD, showing no bound peptide in all three samples. The fourth column shows zoomed in view of the corresponding hexamer NTD, showing clear peptide density in all three samples.

**Figure S5:**
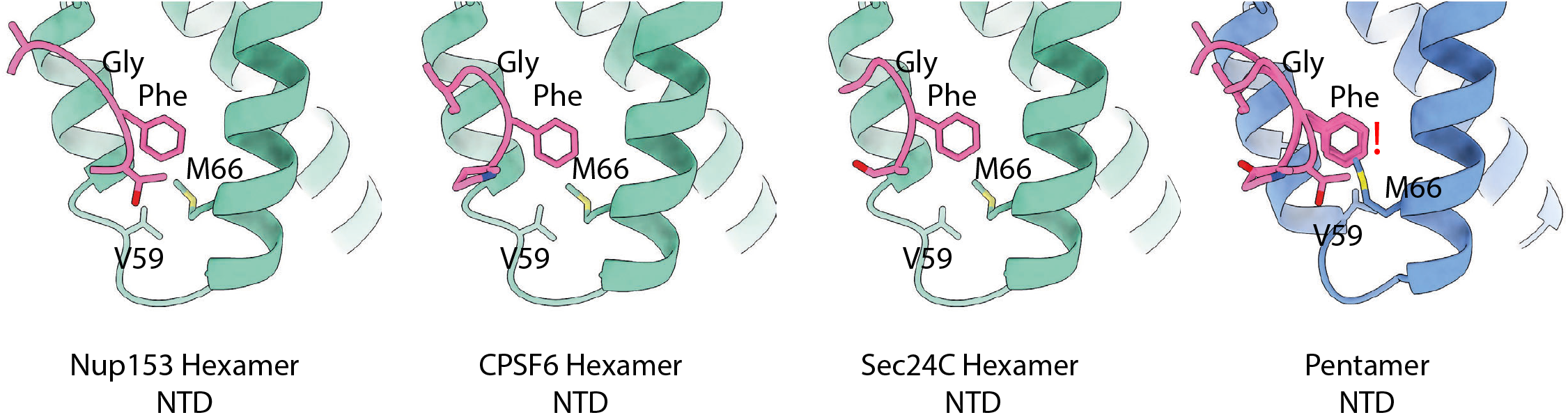
The structure of the pentamer is incompatible with FG repeat motif binding. Models of the FG binding pocket from the CLPs incubated with Nup153_(1407-1429)_, CPSF6_(276-290)_ and Sec24C_(228-242)_ (green), with the bound peptide (pink). On the right the three peptides are superimposed and are shown in the equivalent position in the CA pentamer (blue). The position of M66 in the pentamer is sterically incompatible with binding of the phenylalanine in the FG motif, the position of the clash is denoted by an exclamation mark.

**Figure S6:**
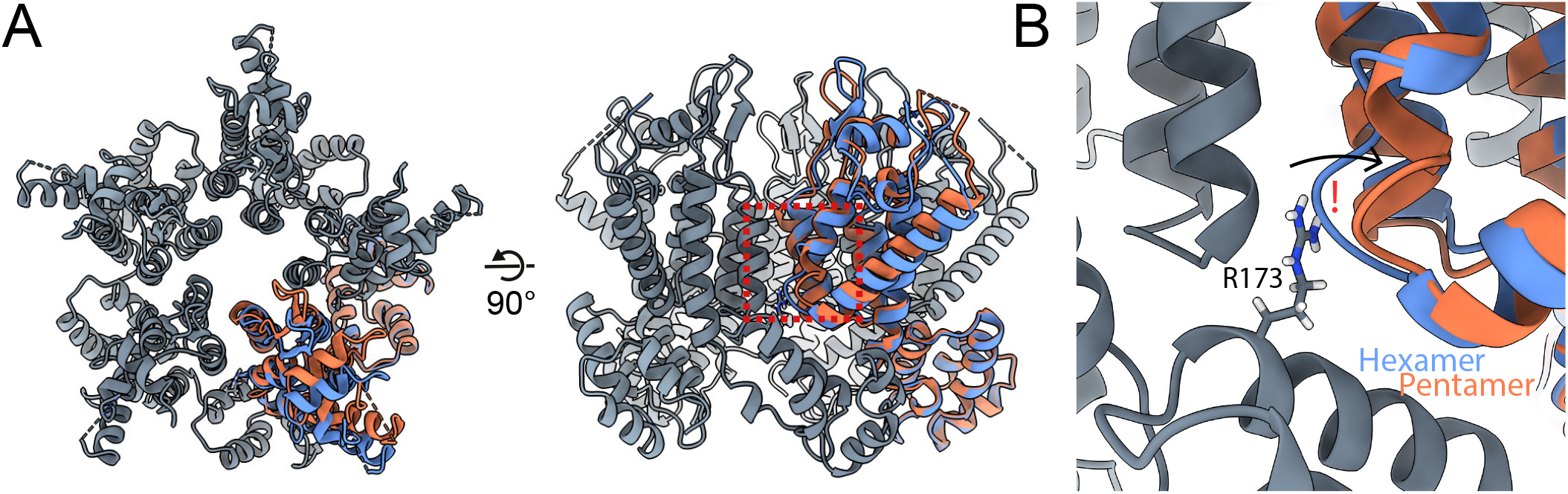
The hexamer-pentamer switch changes to avoid steric clash with R173 in the pentamer. **(A)** A hypothetical pentamer constructed by rigidly fitting CA hexamer monomers into the pentamer reconstruction using only the CTD (grey, one monomer orange). The position of a fit pentamer monomer is also shown for comparison (blue). **(B)** Zoomed in view of the pentamer-hexamer switch region in the theoretical pentamer. R173 of the neighbouring CTD would clash with the pentamer-hexamer switch region if it were not for the conformational change that we observe in the pentamer.

**Table S1.**
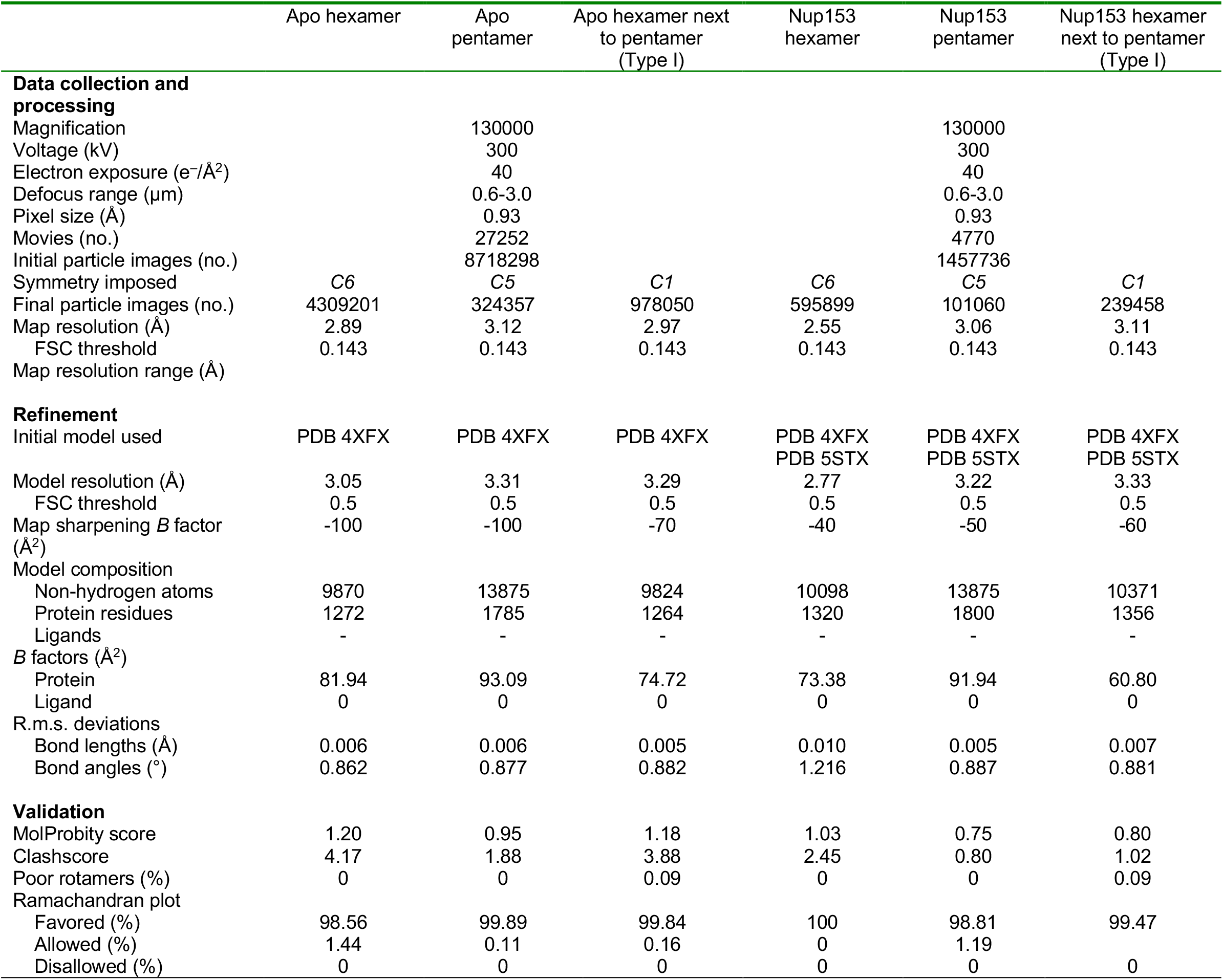
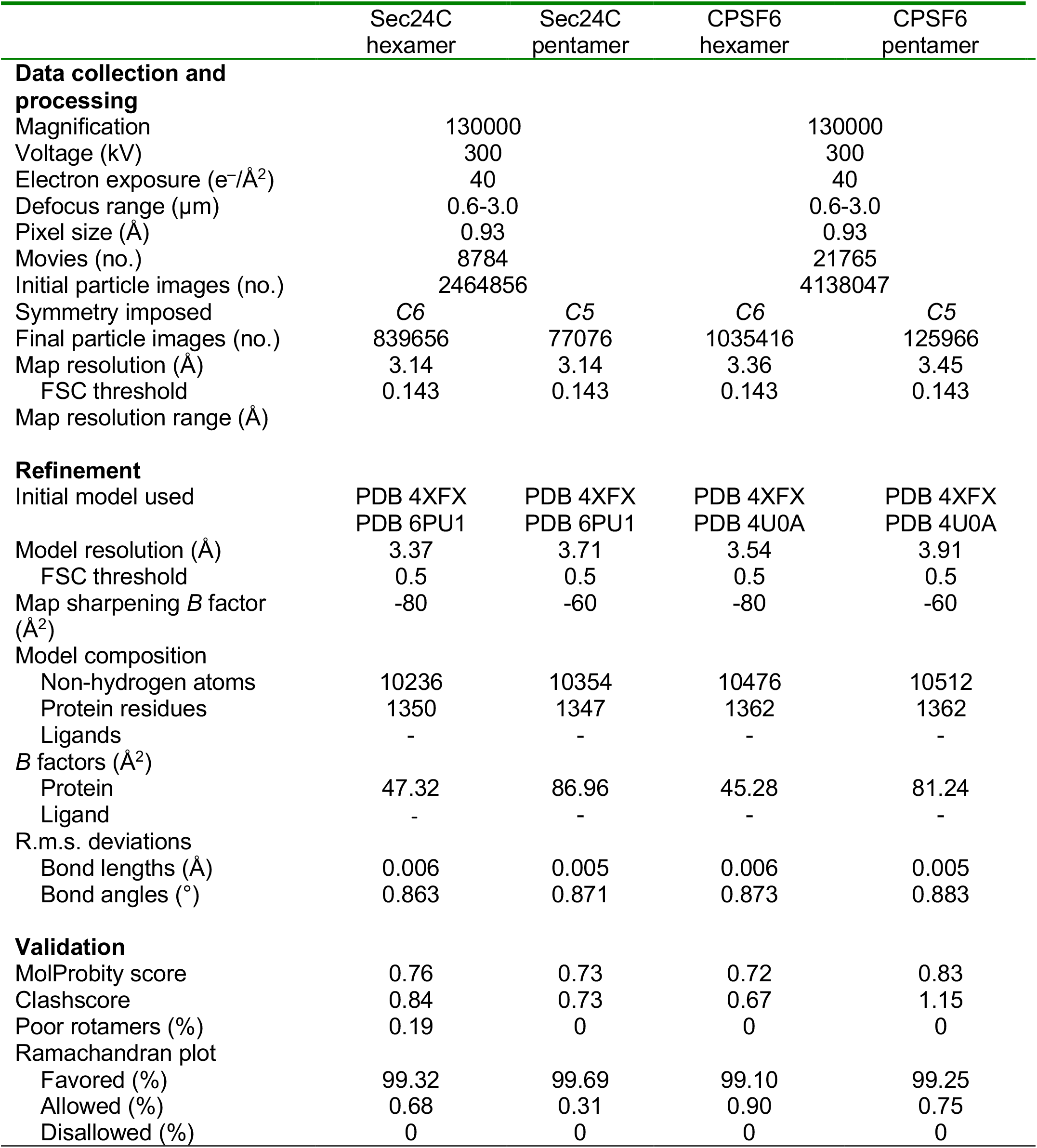
Cryo-EM data collection, refinement and validation statistics

|  | Apo hexamer | Apo pentamer | Apo hexamer next to pentamer (Type I) | Nup153 hexamer | Nup153 pentamer | Nup153 hexamer next to pentamer (Type I) |
| --- | --- | --- | --- | --- | --- | --- |
| **Data collection and processing** |  |  |  |  |  |  |
| Magnification |  | 130000 |  |  | 130000 |  |
| Voltage (kV) |  | 300 |  |  | 300 |  |
| Electron exposure (e<sup>-</sup>/Å<sup>2</sup>) |  | 40 |  |  | 40 |  |
| Defocus range (μm) |  | 0.6-3.0 |  |  | 0.6-3.0 |  |
| Pixel size (Å) |  | 0.93 |  |  | 0.93 |  |
| Movies (no.) |  | 27252 |  |  | 4770 |  |
| Initial particle images (no.) |  | 8718298 |  |  | 1457736 |  |
| Symmetry imposed | C6 | C5 | C1 | C6 | C5 | C1 |
| Final particle images (no.) | 4309201 | 324357 | 978050 | 595899 | 101060 | 239458 |
| Map resolution (Å) | 2.89 | 3.12 | 2.97 | 2.55 | 3.06 | 3.11 |
| FSC threshold | 0.143 | 0.143 | 0.143 | 0.143 | 0.143 | 0.143 |
| Map resolution range (Å) |  |  |  |  |  |  |
| **Refinement** |  |  |  |  |  |  |
| Initial model used | PDB 4XFX | PDB 4XFX | PDB 4XFX | PDB 4XFX<br>PDB 5STX | PDB 4XFX<br>PDB 5STX | PDB 4XFX<br>PDB 5STX |
| Model resolution (Å) | 3.05 | 3.31 | 3.29 | 2.77 | 3.22 | 3.33 |
| FSC threshold | 0.5 | 0.5 | 0.5 | 0.5 | 0.5 | 0.5 |
| Map sharpening *B* factor (Å<sup>2</sup>) | -100 | -100 | -70 | -40 | -50 | -60 |
| Model composition |  |  |  |  |  |  |
| Non-hydrogen atoms | 9870 | 13875 | 9824 | 10098 | 13875 | 10371 |
| Protein residues | 1272 | 1785 | 1264 | 1320 | 1800 | 1356 |
| Ligands | - | - | - | - | - | - |
| *B* factors (Å<sup>2</sup>) |  |  |  |  |  |  |
| Protein | 81.94 | 93.09 | 74.72 | 73.38 | 91.94 | 60.80 |
| Ligand | 0 | 0 | 0 | 0 | 0 | 0 |
| R.m.s. deviations |  |  |  |  |  |  |
| Bond lengths (Å) | 0.006 | 0.006 | 0.005 | 0.010 | 0.005 | 0.007 |
| Bond angles (°) | 0.862 | 0.877 | 0.882 | 1.216 | 0.887 | 0.881 |
| **Validation** |  |  |  |  |  |  |
| MolProbity score | 1.20 | 0.95 | 1.18 | 1.03 | 0.75 | 0.80 |
| Clashscore | 4.17 | 1.88 | 3.88 | 2.45 | 0.80 | 1.02 |
| Poor rotamers (%) | 0 | 0 | 0.09 | 0 | 0 | 0.09 |
| Ramachandran plot |  |  |  |  |  |  |
| Favored (%) | 98.56 | 99.89 | 99.84 | 100 | 98.81 | 99.47 |
| Allowed (%) | 1.44 | 0.11 | 0.16 | 0 | 1.19 |  |
| Disallowed (%) | 0 | 0 | 0 | 0 | 0 | 0 |

|  | Sec24C<br>hexamer | Sec24C<br>pentamer | CPSF6<br>hexamer | CPSF6<br>pentamer |
| --- | --- | --- | --- | --- |
| **Data collection and processing** |  |  |  |  |
| Magnification | 130000 |  | 130000 |  |
| Voltage (kV) | 300 |  | 300 |  |
| Electron exposure (e<sup>-</sup>/Å<sup>2</sup>) | 40 |  | 40 |  |
| Defocus range (μm) | 0.6-3.0 |  | 0.6-3.0 |  |
| Pixel size (Å) | 0.93 |  | 0.93 |  |
| Movies (no.) | 8784 |  | 21765 |  |
| Initial particle images (no.) | 2464856 |  | 4138047 |  |
| Symmetry imposed | C6 | C5 | C6 | C5 |
| Final particle images (no.) | 839656 | 77076 | 1035416 | 125966 |
| Map resolution (Å) | 3.14 | 3.14 | 3.36 | 3.45 |
| FSC threshold | 0.143 | 0.143 | 0.143 | 0.143 |
| Map resolution range (Å) |  |  |  |  |
| **Refinement** |  |  |  |  |
| Initial model used | PDB 4XFX<br>PDB 6PU1 | PDB 4XFX<br>PDB 6PU1 | PDB 4XFX<br>PDB 4U0A | PDB 4XFX<br>PDB 4U0A |
| Model resolution (Å) | 3.37 | 3.71 | 3.54 | 3.91 |
| FSC threshold | 0.5 | 0.5 | 0.5 | 0.5 |
| Map sharpening *B* factor (Å<sup>2</sup>) | -80 | -60 | -80 | -60 |
| Model composition |  |  |  |  |
| Non-hydrogen atoms | 10236 | 10354 | 10476 | 10512 |
| Protein residues | 1350 | 1347 | 1362 | 1362 |
| Ligands | - | - | - | - |
| *B* factors (Å<sup>2</sup>) |  |  |  |  |
| Protein | 47.32 | 86.96 | 45.28 | 81.24 |
| Ligand | - | - | - | - |
| R.m.s. deviations |  |  |  |  |
| Bond lengths (Å) | 0.006 | 0.005 | 0.006 | 0.005 |
| Bond angles (°) | 0.863 | 0.871 | 0.873 | 0.883 |
| **Validation** |  |  |  |  |
| MolProbity score | 0.76 | 0.73 | 0.72 | 0.83 |
| Clashscore | 0.84 | 0.73 | 0.67 | 1.15 |
| Poor rotamers (%) | 0.19 | 0 | 0 | 0 |
| Ramachandran plot |  |  |  |  |
| Favored (%) | 99.32 | 99.69 | 99.10 | 99.25 |
| Allowed (%) | 0.68 | 0.31 | 0.90 | 0.75 |
| Disallowed (%) | 0 | 0 | 0 | 0 |

## Notes

### Competing Interest Statement

The authors have declared no competing interest.

## References

1. S. Mattei, B. Glass, W. J. H. Hagen, H. G. Krausslich, J. A. G. Briggs, The structure and flexibility of conical HIV-1 capsids determined within intact virions. Science 354, 1434–1437 (2016).

2. B. K. Ganser, S. Li, V. Y. Klishko, J. T. Finch, W. I. Sundquist, Assembly and analysis of conical models for the HIV-1 core. Science 283, 80–83 (1999).

3. D. A. Jacques et al., HIV-1 uses dynamic capsid pores to import nucleotides and fuel encapsidated DNA synthesis. Nature 536, 349-+ (2016).

4. X. Lahaye et al., The Capsids of HIV-1 and HIV-2 Determine Immune Detection of the Viral cDNA by the Innate Sensor cGAS in Dendritic Cells. Immunity 39, 1132–1142 (2013).

5. J. Rasaiyaah et al., HIV-1 evades innate immune recognition through specific cofactor recruitment. Nature 503, 402-+ (2013).

6. R. P. Sumner et al., DisruptingHIV-1 capsid formation causescGASsensing of viral DNA. Embo Journal 39 (2020).

7. Q. Shen, C. X. Wu, C. Freniere, T. N. Tripler, Y. Xiong, Nuclear Import of HIV-1. Viruses-Basel 13 (2021).

8. J. Temple, T. N. Tripler, Q. Shen, Y. Xiong, A snapshot of HIV-1 capsid-host interactions. Current Research in Structural Biology 2, 222–228 (2020).

9. T. G. Muller, V. Zila, B. Muller, H. G. Krausslich, Nuclear Capsid Uncoating and Reverse Transcription of HIV-1. Annual Review of Virology 9, 261–284 (2022).

10. R. A. Dick, D. L. Mallery, V. M. Vogt, L. C. James, IP6 Regulation of HIV Capsid Assembly, Stability, and Uncoating. Viruses-Basel 10 (2018).

11. M. Obr, F. K. M. Schur, R. A. Dick, A Structural Perspective of the Role of IP6 in Immature and Mature Retroviral Assembly. Viruses-Basel 13 (2021).

12. N. Renner et al., A lysine ring in HIV capsid pores coordinates IP6 to drive mature capsid assembly. Plos Pathogens 17 (2021).

13. D. L. Mallery et al., IP6 is an HIV pocket factor that prevents capsid collapse and promotes DNA synthesis. Elife 7 (2018).

14. T. Ni et al., Intrinsic curvature of the HIV-1 CA hexamer underlies capsid topology and interaction with cyclophilin A. Nature Structural & Molecular Biology 27, 855-+ (2020).

15. G. P. Zhao et al., Mature HIV-1 capsid structure by cryo-electron microscopy and all-atom molecular dynamics. Nature 497, 643–646 (2013).

16. P. Schommers et al., Changes in HIV-1 Capsid Stability Induced by Common Cytotoxic-T-Lymphocyte-Driven Viral Sequence Mutations. Journal of Virology 90, 7579–7586 (2016).

17. A. T. Gres et al., X-ray crystal structures of native HIV-1 capsid protein reveal conformational variability. Science 349, 99–103 (2015).

18. O. Pornillos, B. K. Ganser-Pornillos, M. Yeager, Atomic-level modelling of the HIV capsid. Nature 469, 424-+ (2011).

19. T. R. Gamble et al., Crystal structure of human cyclophilin A bound to the amino-terminal domain of HIV-1 capsid. Cell 87, 1285–1294 (1996).

20. K. Bichel et al., HIV-1 capsid undergoes coupled binding and isomerization by the nuclear pore protein NUP358. Retrovirology 10 (2013).

21. F. Di Nunzio et al., Human Nucleoporins Promote HIV-1 Docking at the Nuclear Pore, Nuclear Import and Integration. Plos One 7 (2012).

22. A. Bhattacharya et al., Structural basis of HIV-1 capsid recognition by PF74 and CPSF6. Proceedings of the National Academy of Sciences of the United States of America 111, 18625–18630 (2014).

23. A. J. Price et al., Host Cofactors and Pharmacologic Ligands Share an Essential Interface in HIV-1 Capsid That Is Lost upon Disassembly. Plos Pathogens 10 (2014).

24. S. V. Rebensburg et al., Sec24C is an HIV-1 host dependency factor crucial for virus replication. Nature Microbiology 6, 435-+ (2021).

25. Y. Shinkai, M. Kuramochi, T. Miyafusa, New Family Members of FG Repeat Proteins and Their Unexplored Roles During Phase Separation. Frontiers in Cell and Developmental Biology 9 (2021).

26. D. A. Bejarano et al., HIV-1 nuclear import in macrophages is regulated by CPSF6-capsid interactions at the nuclear pore complex. Elife 8 (2019).

27. W. S. Blair et al., New Small-Molecule Inhibitor Class Targeting Human Immunodeficiency Virus Type 1 Virion Maturation. Antimicrobial Agents and Chemotherapy 53, 5080–5087 (2009).

28. K. Singh et al., GS-CA Compounds: First-In-Class HIV-1 Capsid Inhibitors Covering Multiple Grounds. Frontiers in Microbiology 10 (2019).

29. S. Segal-Maurer et al., Capsid Inhibition with Lenacapavir in Multidrug-Resistant HIV-1 Infection. New England Journal of Medicine 386, 1793–1803 (2022).

30. R. A. Dick et al., Inositol phosphates are assembly co-factors for HIV-1. Nature 560, 509-+ (2018).

31. O. Pornillos et al., X-Ray Structures of the Hexameric Building Block of the HIV Capsid. Cell 137, 1282–1292 (2009).

32. Y. F. Chang, S. M. Wang, K. J. Huang, C. T. Wang, Mutations in capsid major homology region affect assembly and membrane affinity of HIV-1 gag. Journal of Molecular Biology 370, 585–597 (2007).

33. U. K. von Schwedler, K. M. Stray, J. E. Garrus, W. I. Sundquist, Functional surfaces of the human immunodeficiency virus type 1 capsid protein. Journal of Virology 77, 5439–5450 (2003).

34. V. Zila et al., Cone-shaped HIV-1 capsids are transported through intact nuclear pores. Cell 184, 1032-+ (2021).

35. S. Q. Zheng et al., MotionCor2: anisotropic correction of beam-induced motion for improved cryo-electron microscopy. Nature Methods 14, 331–332 (2017).

36. D. Kimanius, L. Y. Dong, G. Sharov, T. Nakane, S. H. W. Scheres, New tools for automated cryo-EM single-particle analysis in RELION-4.0. Biochemical Journal 478, 4169–4185 (2021).

37. T. Wagner et al., SPHIRE-crYOLO is a fast and accurate fully automated particle picker for cryo-EM. Communications Biology 2 (2019).

38. A. Punjani, J. L. Rubinstein, D. J. Fleet, M. A. Brubaker, cryoSPARC: algorithms for rapid unsupervised cryo-EM structure determination. Nature Methods 14, 290-+ (2017).

39. A. Punjani, H. W. Zhang, D. J. Fleet, Non-uniform refinement: adaptive regularization improves single-particle cryo-EM reconstruction. Nature Methods 17, 1214-+ (2020).

40. E. F. Pettersen et al., UCSF chimera - A visualization system for exploratory research and analysis. Journal of Computational Chemistry 25, 1605–1612 (2004).

41. E. F. Pettersen et al., UCSF ChimeraX: Structure visualization for researchers, educators, and developers. Protein Science 30, 70–82 (2021).

42. T. I. Croll, ISOLDE: a physically realistic environment for model building into low-resolution electron-density maps. Acta Crystallographica Section D-Structural Biology 74, 519–530 (2018).

43. D. Liebschner et al., Macromolecular structure determination using X-rays, neutrons and electrons: recent developments in Phenix. Acta Crystallographica Section D-Structural Biology 75, 861–877 (2019).

44. V. B. Chen et al., MolProbity: all-atom structure validation for macromolecular crystallography. Acta Crystallographica Section D-Structural Biology 66, 12–21 (2010).

45. A. Punjani, D. J. Fleet, 3D variability analysis: Resolving continuous flexibility and discrete heterogeneity from single particle cryo-EM. Journal of Structural Biology 213 (2021).

